# Immunologically mediated trade-offs shaping transmission of sylvatic dengue and Zika viruses in native and novel non-human primate hosts

**DOI:** 10.1101/2023.06.30.547187

**Authors:** Kathryn A. Hanley, Hélène Cecilia, Sasha R. Azar, Brett Moehn, Wanqin Yu, Ruimei Yun, Benjamin M. Althouse, Nikos Vasilakis, Shannan L. Rossi

## Abstract

Mosquito-borne dengue (DENV) and Zika (ZIKV) viruses originated in Old World sylvatic cycles involving monkey hosts, spilled over into human transmission, and were translocated to the Americas, creating potential for spillback into neotropical sylvatic cycles. Studies of the trade-offs that shape within-host dynamics and transmission of these viruses are lacking, hampering efforts to predict spillover and spillback. We exposed native (cynomolgus macaque) or novel (squirrel monkey) hosts to mosquitoes infected with either sylvatic DENV or ZIKV and monitored viremia, natural killer cells, transmission to mosquitoes, cytokines, and neutralizing antibody titers. Unexpectedly, DENV transmission from both host species occurred only when serum viremia was undetectable or near the limit of detection. ZIKV replicated in squirrel monkeys to much higher titers than DENV and was transmitted more efficiently but stimulated lower neutralizing antibody titers. Increasing ZIKV viremia led to greater instantaneous transmission and shorter duration of infection, consistent with a replication-clearance trade-off.

## Introduction

Spillover of zoonotic, arthropod-borne viruses (arboviruses) into humans is accelerating, with outcomes ranging from dead-end infections to pandemics ^1^. Because of the unparalleled mobility of humans, these viruses are being transported across the world, creating the potential for spillback into novel wildlife reservoirs ^2^. Such translocation is not new: yellow fever virus (YFV), which originated in a sylvatic cycle in Africa involving non-human primates (NHPs) and arboreal mosquitoes was carried via sailing ships to the neotropics centuries ago ^3,4^. Soon after, YFV established urban transmission in the Americas and spilled back into a sylvatic cycle maintained in neotropical primates and mosquitoes that plagues South America to present day ^3,4^. The four serotypes of dengue virus (DENV-1-4) underwent a similar journey, albeit their ancestral sylvatic cycles occur in Asia, but DENV has never spilled back into a sylvatic reser - voir in the Americas ^5^. Thus, when ZIKV was detected in the Americas in 2015, there was great concern but equally great uncertainty, about its potential to establish a neotropical sylvatic cycle ^6-8^. While multiple instances of ZIKV infection in neotropical primates have now been documented ^9,10^, the potential for such spillback events to launch a sylvatic cycle remains unclear. Furthermore, the sylvatic cycles of DENV and ZIKV persist in Asia and Africa, where both continue to spill over into humans, e.g. ^11-17^.

Mathematical models using within-host viral dynamics to predict between-host transmission are key for predicting arbovirus spillover, spread, and spillback ^18^. These models assume within-host trade-offs that influence virus transmission ^19^, particularly the trade-off between instantaneous pathogen transmission and infection duration ^20,21^. Host mortality is often invoked in theoretical studies as the mechanism that regulates infection duration, however most pathogens are cleared by the immune response. Indeed, for sylvatic DENV and ZIKV there is no evidence of disease in infected Old World monkeys ^22-24^, although sylvatic ZIKV was first isolated from a febrile sentinel macaque in Uganda ^14^ and poses similar risks to the developing fetus in Old World monkeys as do human-endemic lineages of the virus ^25,26^. Thus, transmission of these viruses likely depends on the trade-off between the magnitude of viral replication and immune clearance ^27^.

Studies of replication-clearance trade-offs in arboviruses are scarce. The majority of studies on the intrahost dynamics of arboviruses during experimental infection of vertebrate reservoir hosts found an inverse relationship between the magnitude (peak titer) of infection and duration of infection ^28,29^. However, most of these studies delivered virus to the host via needle injection, which differs from natural transmission via mosquito bite ^30-32^. We formulated a dynamical model comparing the effect of a “tortoise” strategy of low magnitude, long-duration viremia and a “hare” strategy of short-duration, high-magnitude viremia and showed that arboviruses adopting a tortoise strategy had higher rates of persistence in both host and vector populations ^29^. Ben-Shachar and Koelle ^33^ investigated the replication-clearance trade-off in DENV using within-host simulation models based on data from a human cohort study in which mosquitoes fed on infected individuals and found that a replication-clearance trade-off selects for intermediate DENV virulence. Importantly for the current study, they invoked natural killer (NK) cells as the mechanism of innate immunity driving this effect.

In the current study, we investigated trade-offs between replication and clearance of sylvatic DENV and ZIKV, as well as immune responses driving these trade-offs, in both native and novel NHP hosts. This study tested three *a priori* hypotheses: Hypothesis 1 - sylvatic arboviruses experience a replication-clear - ance trade-off in both native and novel hosts, as evidenced by a positive relationship between the dose of virus delivered with peak virus titer, a positive relationship between virus titer and transmission to mos - quitoes, and a negative relationship between peak virus titer and duration of viremia; Hypothesis 2 - NK cells shape this trade-off, as evidenced by a negative relationship between NK cell mobilization immediately post-infection (pi) and peak virus titer as well as levels of neutralizing antibody, and Hypothesis 3 - sylvatic arboviruses experience different replication-clearance trade-offs in native hosts and novel hosts, resulting in less transmission from novel hosts.

## Results

### Overview of experimental design

The native host in this study was the cynomolgus macaque, *Macaca fascicularis* ^34-36^, and the novel host was the neotropical squirrel monkey, *Saimiri boliviensis boliviensis*. Virus was delivered by infected *Ae. albopictus* mosquitoes, which are native to Asia but distributed across much of the world, a likely bridge vector for spillover and spillback of both DENV and ZIKV ^37^, and a vector in outbreaks of sylvatic ZIKV in Gabon ^38^.

Figure 1. shows the experimental design for this study. To infect macaques, cartons of either 1 (low dose, n = 5) or 10 (high dose, n = 5) *Ae. albopictus* that had been intrathoracically (IT) inoculated with a sylvatic Malaysian strain of DENV-2, or 10 uninoculated, control *Ae. albopictus* (n = 3) were placed upon a monkey’s ear for approximately 10 minutes. Engorged mosquitoes were separated, incubated for 2 days, and force-salivated to estimate dose of virus delivered. Squirrel monkeys were infected with either a high dose (15 mosquitoes) of sylvatic DENV-2 (n = 10 monkeys) or a sylvatic African ZIKV strain (n = 10) or control mosquitoes (n = 4). Because the small size of squirrel monkeys constrained blood draw frequency, monkeys were assigned to one of two cohorts and bled prior to infection and on alternating days in the first week pi. Cytokines, NK cells, neutralizing antibody via plaque neutralization reduction titers (PRNTs), weight and temperature were measured on designated days prior to and post-infection (dpi). On designated dpi, uninfected *Ae. albopictus* were allowed to feed on each animal to monitor transmission. All raw data from macaques infected with DENV-2 and squirrel monkeys infected with DENV-2 or ZIKV, save for temperature and PRNT values against non-infecting viruses, are provided in **Supplemental Tables S.1, S.2** and **S.3**, respectively.

**Figure 1.**
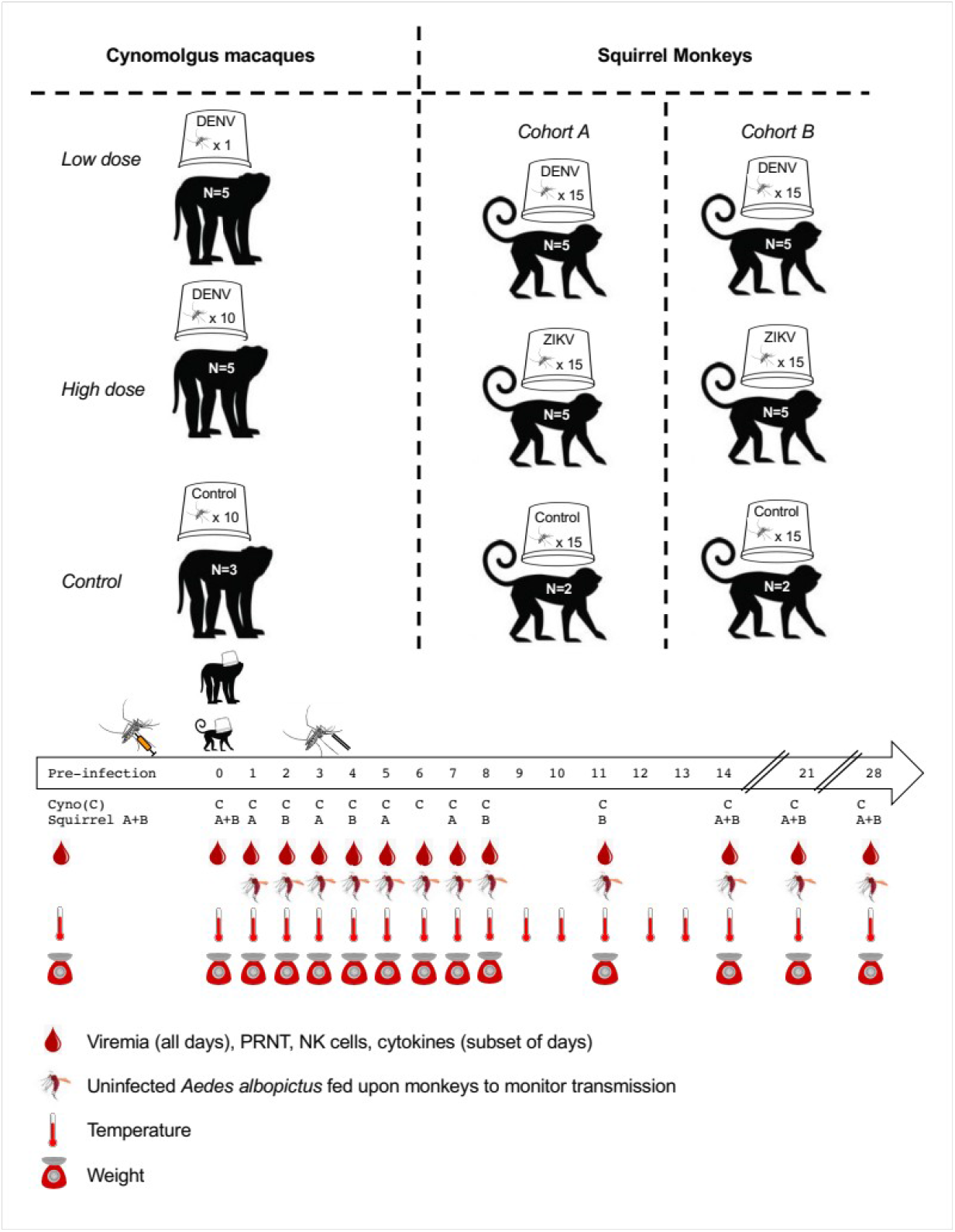
Overview of experimental infections of cynomolgus macaques and squirrel monkeys with DENV-2 or ZIKV and subsequent sampling. Blood samples were used to measure viremia, NK cells, and neutralizing antibody. Monkey body temperature was monitored continuously via implanted transponder and weight was monitored on designated days. Infection and monitoring were similar in macaques and squirrel monkeys, save that: (i) the sample size of infected and uninfected mosquitoes was increased from 10 in macaques to 15 in squirrel monkeys, (ii) stool was collected daily from macaques to screen for occult blood; stool was not collected from squirrel monkeys due to pair housing of animals, (iii) mosquitoes fed on squirrel monkeys on dpi 3 and 4 were force-salivated after a 14 day incubation, but this was not done in macaques, and (iv) squirrel monkeys were euthanized at the conclusion of the experiment and necropsies were conducted; euthanasia was not performed on macaques. In one instance, blood sampling (dpi 7) and mosquito feeding (dpi 8) occurred on different days for macaques.

### Sylvatic DENV-2 replicated to low levels in macaques, stimulating high neutralizing antibody titers but resulting in no signs or symptoms of infection

DENV-2 was not initially detectable in the saliva of any of the IT inoculated mosquitoes used to deliver virus to macaques, but after one passage, virus was detected in a subset of salivations (**Table 1**). Thus, we used number of mosquitoes salivating detectable virus to estimate total dose of virus delivered. For macaque UG253A, the single mosquito that fed produced no detectable virus in the saliva, and UG253A never produced detectable viremia or seroconverted (defined here as a PRNT50 titer, the most permissive cutoff, above the limit of detection (LOD), **Supplemental Table S.4**). Hence UG253A was reassigned, *post hoc*, to the control treatment. Macaque SB393 received a single bite from a mosquito that did not salivate detectable virus, but this monkey seroconverted. Of remaining 8 monkeys in the low and high dose treatments, 7 seroconverted (**Table 1**). All statistical analyses of neutralizing antibody levels were conducted using PRNT80 values, a more stringent measure consistent with our previous study of sylvatic DENV-2 in African green monkeys ^22^. The harmonic mean PRNT80 value (values < LOD set at 19) for macaques was 69.7. Viremia was detected from serum in only three macaques, one from the low dose and two from the high dose treatment (**Table 1** and **Fig. 2A**).

**Table 1:** Assignment of each cynomolgus macaque to high dose (10 DENV-2 infected mosquitoes), low dose (1 DENV-2 infected mosquito) or control (10 uninfected mosquitoes) treatments and subsequent viremia, transmission to mosquitoes, and neutralizing antibody response.

| NHP ID | Sex | *A priori*<br>mosquito<br>treatment | *Post hoc*<br>mosquito<br>Treatment<sup>1</sup> | No. infected<br>mosquitoes<br>engorged | No. infected<br>mosquitoes<br>with de-<br>tectable<br>virus in<br>saliva post<br>passage<sup>2</sup> | Days on<br>which virus<br>was de-<br>tectable in<br>serum, <u>day of</u><br><u>peak titer</u><br>(log<sub>10</sub>pfu/ ml) | Days on<br>which virus<br>was trans-<br>mitted to<br>mosquitoes<br>(% in-<br>fected<sup>4</sup>) | PRNT80 on<br>Day |  |
| --- | --- | --- | --- | --- | --- | --- | --- | --- | --- |
|  |  |  |  |  |  |  |  | -10 | 28 |
| NV289 | M | Control | Control | 8 | 0 | - | - | <20 | <20 |
| UG171 | F | Control | Control | 5 | 0 | - | - | <20 | <20 |
| NV259 | F | Control | Control | 9 | 0 | - | - | <20 | <20 |
| SB393 | M | Low Dose | Low Dose | 1 | 0 | - | - | <20 | 40 |
| FR469A | M | Low Dose | Low Dose | 1 | 1 | 4 (3.9) | - | <20 | 160 |
| UG253A<sup>1</sup> | M | Low Dose | Control | 1 | 0 | - | - | <20 | <20 |
| BC407 | F | Low Dose | Low Dose | 1 | 1 | - | - | <20 | <20 |
| CP60 | F | Low Dose | Low Dose | 1 | 1 | - | - | <20 | 80 |
| BC116A | M | High Dose | High Dose | 7 | 6 | 4 (na<sup>3</sup>) | - | <20 | 40 |
| FR423A | M | High Dose | High Dose | 8 | 5 | 4, 7 (2.5) | 5 (60%) | <20 | 320 |
| SB395 | F | High Dose | High Dose | 8 | 3 | - | - | <20 | 1280 |
| FR840 | F | High Dose | High Dose | 9 | 4 | - | 8 (100%) | <20 | 320 |
| BC167 | F | High Dose | High Dose | 7 | 3 | - | 5 (20%) | <20 | 1280 |

1. See text for justification of change from *a priori* to *post-hoc* assignment 2. Includes only mosquitoes that survived the two-day rest period post-feeding. Survival was generally very high; survival data is available in Supplemental Table S.1 3. Not applicable as viremia was detected only post-passage 4. Considered positive if virus detected in bodies or legs, which were processed separately

**Figure 2.**
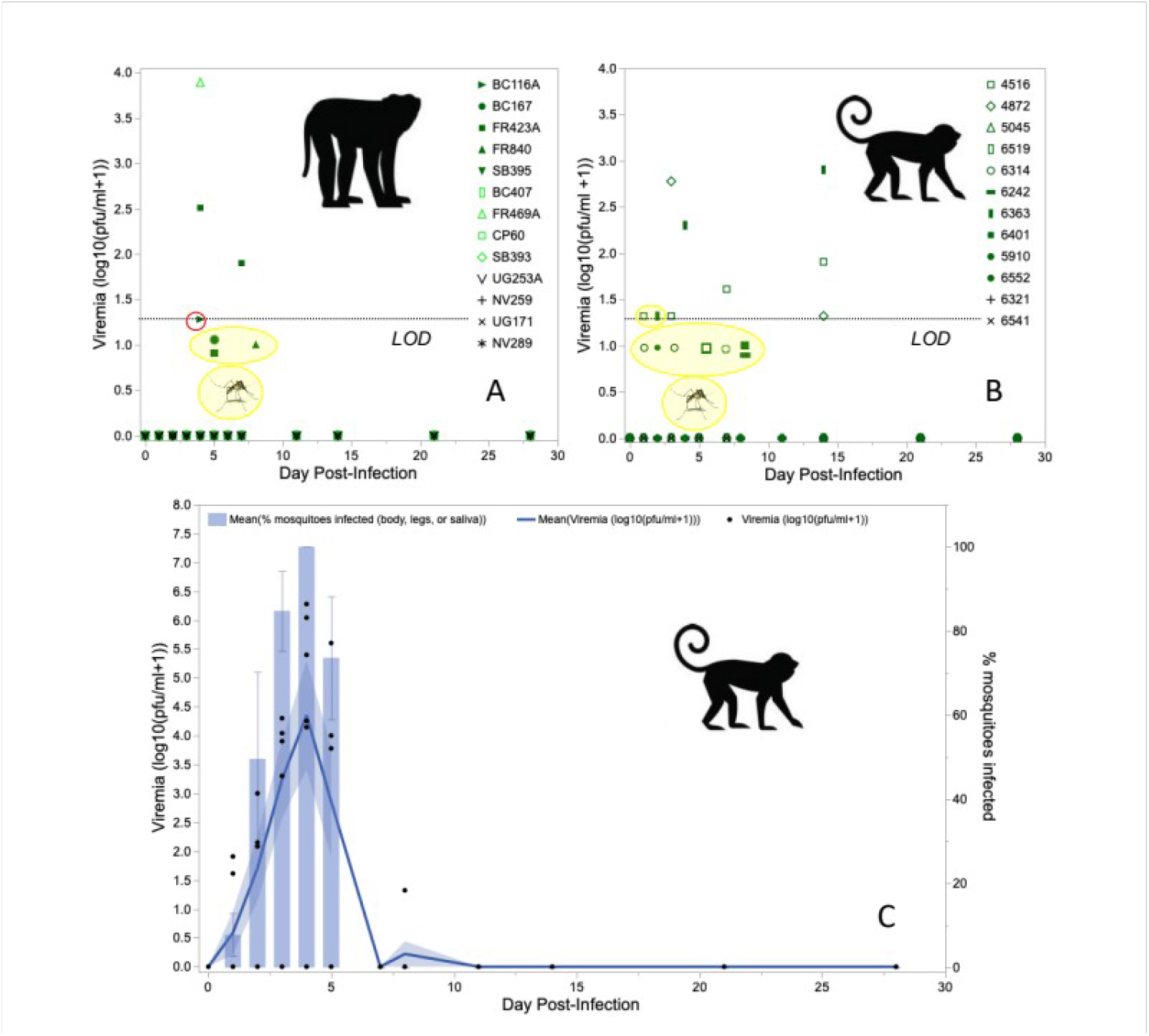
Replication and transmission to *Ae. albopictus* of: A) sylvatic DENV-2 in cynomolgus macaques, B) sylvatic DENV-2 in squirrel monkeys, C) sylvatic ZIKV in squirrel monkeys. In panel A, the red circle shows detection of viremia only after passage of serum (and therefore below the LOD). Light shaded symbols indicate low dose of virus (1 mosquito), dark shaded symbols indicate high dose (10-15 mosquitoes), and black symbols indicate controls (10-15 uninfected mosquitoes). In panels B and C, filled symbols indicate cohort 1 and open symbols indicate cohort 2, per the experimental design in Figure 1. In panels A and B, yellow circles and highlight indicate detection of transmission to mosquitoes; when these values fall below the LOD, there was no detection of viremia in monkey sera through direct titration or passage. Y axis range differs for DENV-2 (panels A and B) and ZIKV (panel C). Viremia was monitored by direct titration of serum as well as one passage of serum followed by titration; transmission was monitored by feeding cartons of 10-15 uninfected mosquitoes on each monkey. Error bands and bars indicate 1 SE.

Macaque body temperatures stayed within the normal range for this species ^39,40^ and there were no differences among treatment groups (**Fig. S.1A**). There were no differences among groups in changes in body weight following treatment (**Fig. S.2A**). Occult blood was detected in stool only once, in a control monkey. Cytokines showed substantial inter-individual variation with no sex differences at baseline (**Supplemental text S.1 Table S.5.1**), no long-term disruption of the cytokine equilibrium (comparison of baseline and day 28, **Supplemental text S.2, Table S.5.2**), and no strong effects of treatment (**Supplemental text S.3.1, Table S.5.3**).

### Macaques were more likely to become infected as dose of DENV-2 increased, but transmission to mosquitoes only occurred when viremia was not detectable from serum

Only three macaques transmitted DENV-2 to mosquitoes (**Table 1, Fig. 2A, Supplemental Table S.6)**. Intriguingly, DENV-2 was not detected from raw or passaged serum from any monkey on the day of transmission or, in the case of monkeys FR840 and BC167, at any point during the study. Too few monkeys produced detectable virus in serum to analyze peak virus titer or duration of infection, but it was possible to analyze the association of total dose on the likelihood of becoming viremic, as evidenced by detectable virus in raw or passaged serum or infection of mosquitoes (n = 5 viremic macaques total). Consistent with Hypothesis 1, detectably viremic animals received a significantly greater number of infectious bites than animals that were not detectably viremic (logistic regression, DF = 1, n = 9, χ^2^ = 4.63, P = 0.03).

### DENV-2 dose did not affect NK cell mobilization or neutralizing antibody titers in macaques

To analyze variation in NK cells, we used the percent NK cells of the total number of peripheral blood mononuclear cells (PBMCs). After Bonferroni correction for multiple testing, values for % NK cells in individual, infected macaques at each dpi were significantly correlated with the following dpi, however values at D0 and D1 dpi were not significantly correlated. Thus, we analyzed only % NK cells from 1 dpi. Day 1 % NK cells were not significantly different among control, low dose and high dose macaques (ANOVA, DF = 2, F =2.74, P = 0.11) (**Fig. 3A**), and Day 28 PRNT80 values were not significantly different between high and low dose macaques (t-test on log_10_ transformed data, DF = 7, t = 2.29, P = 0.056) (**Table 1**). Consistent with Hypothesis 2, there was a negative relationship in infected macaques between dpi 1 % NK cells and PRNT80 values (log_10_ PRNT80 decreased by 0.12 (95% CI (−0.25; 0.003)) for every 1% increase in %NK cells), albeit the difference was not significant (DF = 1, F = 5.3, P = 0.055, adjusted R^2^ = 0.35) (**Fig. 3D**). % NK cells from the first week post-infection showed no significant association with IL-12 concentrations in control or infected macaques (**Supplemental text S.4.2, Fig. S.3, S.4**).

**Figure 3.**
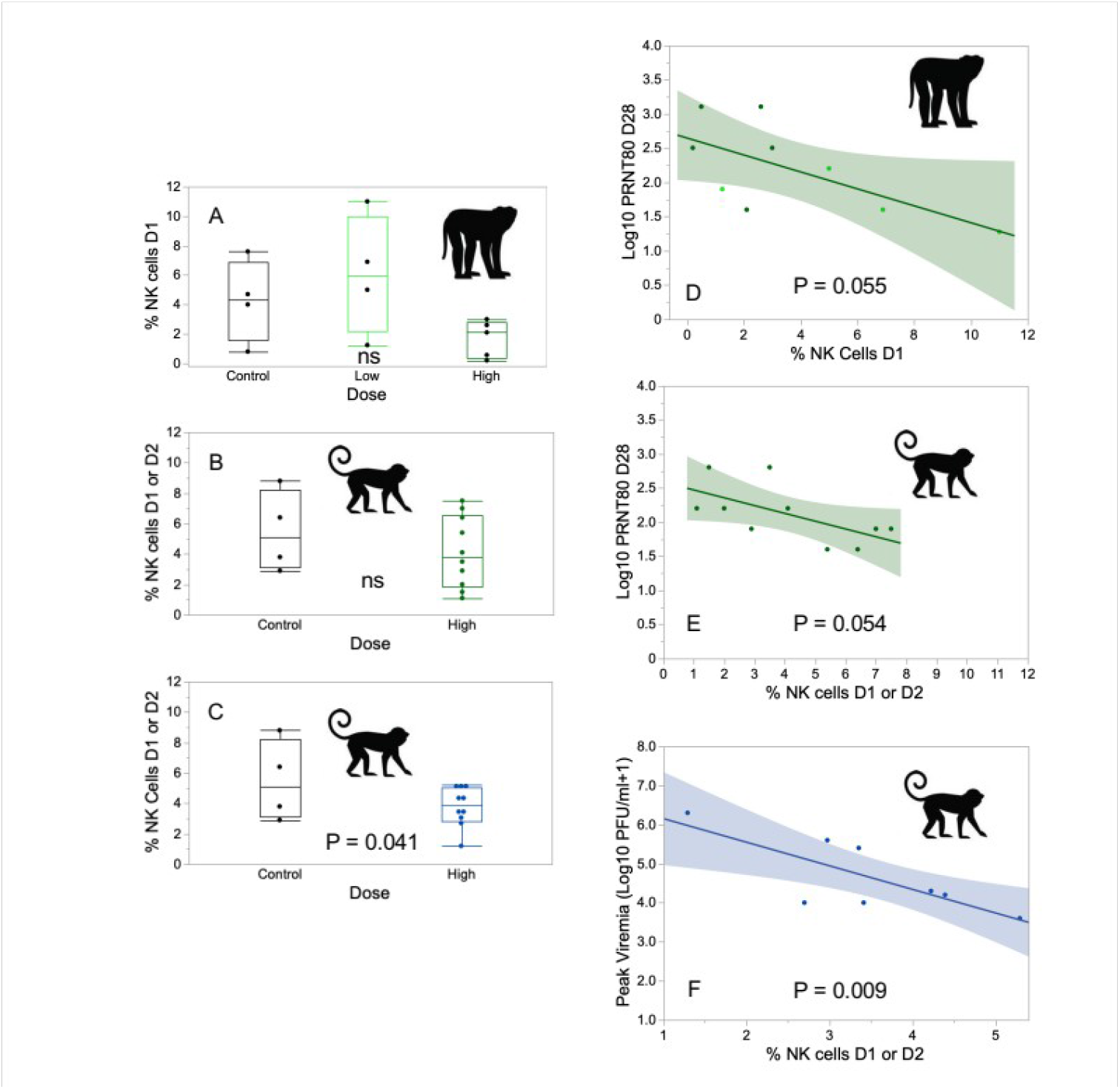
Effect of treatments on early % NK cells (A-C), associations between early % NK cells and PRNT80 (D-E) or peak virus titer (F): A) % NK cells on dpi 1 in cynomologus macaques infected with control, low dose or high dose DENV-2; B) % NK cells on dpi 1 or 2 in squirrel monkey infected with control or high dose DENV; C) % NK cells on dpi 1 or 2 in squirrel monkey infected with control or high dose ZIKV (note that the same 4 control monkeys are shown in panels C and D); D) regression of log_10_ transformed PRNT80 values on % NK values on dpi 1 in cynomolgus macaques infected with low (light green symbols) or high dose (dark green symbols) DENV (controls are excluded from this analysis); E) regression of log_10_ transformed PRNT80 values on dpi 28 against a % NK cells on dpi 1 or 2 in squirrel monkeys infected with high dose DENV-2 (controls are excluded from this analysis); F) regression of peak virus titer on % NK cells on dpi 1 or 2 in squirrel monkeys infected with high dose ZIKV (controls are excluded from this analysis). Note the difference in X axis between panel F versus D and E.

### Sylvatic DENV-2 replicated to low levels in squirrel monkeys, stimulating high neutralizing antibody titers but resulting in no signs or symptoms of infection

DENV-2 was detectable in the saliva of 30 IT inoculated mosquitoes by titer of mosquitoes or passaged homogenates (**Table 2**). All monkeys in the high dose treatment seroconverted (PRNT50> LOD), with a harmonic mean PRNT80 value of 91.4, but only three produced detectable viremia in serum (**Table 2, Fig. 2B**). Body temperatures for infected squirrel monkeys stayed within the normal range for this species and there were no differences between treatment groups (**Supplemental Fig. S.1B)**. There were no differences between infected and control monkeys in body weight change (**Supplemental Fig. S.2B**). No sex differences at baseline (**Supplemental text S.1, Table S5.1**), nor long-term disruption of the cytokine equilibrium (comparison of baseline and day 28, **Supplemental text S.2, Table S.5.2**), nor strong effects of infection (**Supplemental text S.3.1, Table S.5.3**) on cytokine concentrations were apparent. There were no notable findings in necropsy reports on these animals.

**Table 2:** Virus delivered to squirrel monkeys via cartons of 15 mosquitoes and subsequent viremia, transmission to mosquitoes, and neutralizing antibody response.

| NHP ID | Sex | Virus treatment | No. infected mosquitoes engorged | No. infected mosquitoes salivating virus <sup>1</sup> | Total dose of virus delivered (log<sub>10</sub> pfu) | Days on which virus was detectable in serum, day of peak titer, (peak titer (log<sub>10</sub>pfu/ ml)) | Days on which virus was transmitted to mosquitoes (% infected<sup>4</sup>) | PRNT80 on Day |  |
| --- | --- | --- | --- | --- | --- | --- | --- | --- | --- |
|  |  |  |  |  |  |  |  | -7 | 28 |
| 6314 | M | DENV | 3 | 2 | 0.3 | - | 1 (11%)<br>3 (8%)<br>7 (38%) | <20 | 640 |
| 6519 | M | DENV | 4 | 4 | 1.6 | - | - | <20 | 40 |
| 4516 | F | DENV | 5 | 4 | 3.2 | 1, 3, 7, **14** (1.9) | 1(100%)<br>5 (13%) | <20 | 640 |
| 5045 | F | DENV | 4 | 2 | 3.3 | - | - | <20 | 80 |
| 4872 | F | DENV | 3 | 3 | 1.6 | **3** (2.8), 14 | - | <20 | 160 |
| 6401 | M | DENV | 4 | 2 | 1.6 | - | 8 (22%) | <20 | 40 |
| 6363 | M | DENV | 5 | 2 | 0.3 | 2, 4, **14** (2.9) | 2 (44%) | <20 | 80 |
| 6552 | M | DENV | 4 | 3 | 2.3 | - | 2 (40%) | <20 | 80 |
| 6242 | F | DENV | 4 | 3 | 2.3 | - | 8 (57%) | <20 | 160 |
| 5910 | F | DENV | 6 | 5 | 2.3 | - | - | <20 | 160 |
| 6541 | M | Control | 2 | 0 | 0 | - | - | <20 | <20 |
| 6321 | M | Control | 4 | 0 | 0 | - | - | <20 | <20 |
| 6550 | M | ZIKV | 9 | 6 | 3.5 | 1, **3**(4.0), 5 | 3 (35%)<br>5 (46%) | <20 | 80 |
| 6518 | M | ZIKV | 5 | 1 | 2.8 | **3** (3.3) | 1 (14%)<br>3 (14%)<br>5 (10%) | <20 | <20 |
| 6311 | M | ZIKV | 10 | 5 | 3.7 | **3** (3.9), 5 | 3 (50%)<br>5 (9%) | <20 | <20 |
| 5013 | F | ZIKV | 11 | 11 | 3.9 | 1, 3, **5** (4.0) | 3 (22%)<br>5 (71%) | <20 | <20 |
| 4806 | F | ZIKV | 4 | 4 | 2.9 | 3, **5** (5.6) | **5** (64%) <sup>2</sup> | <20 | <20 |
| 6347 | M | ZIKV | 6 | 6 | 3.2 | **4** (5.4) | 2 (17%)<br>4 (80%) | <20 | 40 |
| 6359 | M | ZIKV | 7 | 7 | 4.2 | 2, **4** (4.2) | 4 (60%) | <20 | <20 |
| 5730 | F | ZIKV | 4 | 4 | 3.1 | 2, **4** (4.3) | 4 (100%) | <20 | 40 |
| 4683 | F | ZIKV | 7 | 6 | 3.5 | 2, **4** (6.0), 8 | 2 (50%)<br>4 (25%) | <20 | na<sup>3</sup> |
| 4728 | F | ZIKV | 9 | 8 | 3.9 | 2, **4** (6.3) | 4 (50%) | <20 | na<sup>3</sup> |
| 5929 | F | Control | 8 | 0 | 0 | - | - | <20 | <20 |
| 5080 | F | Control | 10 | 0 | 0 | - | - | <20 | <20 |

1. Includes only mosquitoes that survived the two-day rest period post-feeding. Survival was generally very high; survival data is available in Supplemental Table S.1. When virus was only detectable after passage, the dose was arbitrarily estimated at 1 pfu per salivation
2. No mosquitoes that fed on Day 3 on this monkey survived incubation
3. Euthanized before Day 28
4. Considered positive if virus detected in bodies, legs, or saliva, which were processed separately

### Squirrel monkeys were not more likely to become infected as dose of DENV-2 increased, and transmission to mosquitoes only occurred when viremia was not detectable from serum or was slightly higher than the LOD

Three monkeys produced viremia that could be detected from serum, and, as with macaques, transmission to mosquitoes occurred when viremia was undetectable or just above the LOD. Two monkeys that produced viremia detectable in serum infected mosquitoes, but only when their viremia was at its lowest point. Another four monkeys that did not produce detectable viremia nonetheless infected mosquitoes (**Table 2 and Fig. 2B**). Contra Hypothesis 1, dose of DENV delivered had no significant impact on the likelihood of squirrel monkeys becoming detectably viremic (logistic regression, N = 7, Table 2, DF = 1, χ^2^ = 1.0556, P = 0.30). DENV-2 was not detectable in any of the salivations from the mosquitoes that fed on monkeys at 3 and 4 dpi (**Supplemental Table S.6**). Given the small number of monkeys for which peak titer could be quantified, we did not analyze the associations with peak titer.

### In squirrel monkeys infected with DENV-2, NK cells mobilized early in infection were negatively associated with neutralizing antibody titers

Early % NK cells (1 or 2 dpi) in infected squirrel monkeys did not differ from controls (t-test, DF = 12, t = 0.93, P = 0.37; note that all four control squirrel monkeys were used in comparisons with both DENV and ZIKV infected monkeys; **Fig. 3B**). Percent NK cells in infected monkeys at early (1 or 2 dpi) and later (7 or 8 dpi) timepoints were significantly, positively correlated with each other (P = 0.005); early % NK cells were correlated with % NK cells prior to infection (P = 0.04), but later % NK cells were not (P = 0.21). We did not analyze the association of early % NK cells and peak virus titer. Similar to DENV-2 in cynomolgus macaques, and in line with Hypothesis 2, there was a negative relationship (log_10_ PRNT80 decrease by 0.11 95%CI (−0.23; 0.002) for every 1% increase in %NK cells) in infected monkeys between early % NK cells and PRNT80 values on Day 28 (**Fig. 3E**), albeit the difference was not significant (DF = 1, F = 5.1, P = 0.054, adjusted R^2^ = 0.31).

### Sylvatic DENV-2 dynamics were similar in native and novel hosts

Squirrel monkeys and macaques received similar doses of DENV-2 (**Supplemental text S.4.1, S.4.2, S.4.3**). The only significant effect on PRNT80 values revealed by a linear model testing the effects of early % NK cells and host species, as well as the interaction between the two factors, was early % NK cells (DF= 1, F = 10.8, P = 0.005). Eotaxin, I-TAC, and RANTES concentrations were higher in infected macaques than infected squirrel monkeys (**Supplemental text S.5, Table S.5.4, Fig. S.5**); this difference was attributable to species identity rather than infection, as the species differences for these cytokines were of the same magnitude in control monkeys (**Table S.5.4, Fig. S.5**). Transmission from both macaques and squirrel monkeys was too infrequent for statistical comparison, but provided no support for Hypothesis 3, that transmission is lower from a novel host. Intriguingly, DENV transmission from both macaques and squirrel monkeys was associated with low or undetectable viremia (**Fig. 2A,B**).

### Sylvatic ZIKV replicated to high levels in squirrel monkeys, stimulating low neutralizing antibody titers

ZIKV was detectable in the saliva of 58 intrathoracically inoculated mosquitoes. There was a tight relationship between number of mosquitoes salivating ZIKV and total dose delivered (DF = 1, F = 12.6, P = 0.008, adjusted R^2^ = 0.561, slope 0.14 (0.05 ; 0.22) in log_10_ units). All monkeys receiving bites from ZIKV-infected mosquitoes produced detectable viremia and infected mosquitoes on at least one dpi and all seroconverted (**Table 2, Fig. 2C, Table S.4, Fig. S.6**) with a mean harmonic PRNT80 of 23.6. ZIKV titer peaked between dpi 3 and 5 and viremia lasted between 3 to 7 days, although there is substantial uncertainty to these estimates given the sampling regimen.

### Squirrel monkeys infected with ZIKV showed lower levels of RANTES than controls, and two monkeys showed significant pathology that was inconsistent with that previously documented for ZIKV

Body temperatures for squirrel monkeys infected with ZIKV stayed within the normal range for this species and there were no differences among treatment groups (**Fig. S.1C**). There were no differences between infected and control monkeys in body weight change, and no monkeys dipped more than 10% in body weight (**Fig S.2C**). No sex differences at baseline (**Supplemental text S.1, Table S5.1**), nor longterm disruption of the cytokine equilibrium (comparison of baseline and day 28, **Supplemental text S.2, Table S.5.2**) were apparent. Among all cytokines analyzed, the concentration of RANTES was significantly higher in controls than in ZIKV-infected squirrel monkeys (difference of 0.27 (0.14 ; 0.41) log_10_ pg/μl, P = 8.82e-5, **Supplemental Text S.3, Table S.5.3, Fig. S.7**). As described in **Supplemental Text S.6**, two monkeys infected with ZIKV had to be euthanized prior to the end of the experiment following the recommendation of the head veterinarian on staff: 4683 due to cluster seizures and 4728 due to a worsening sore on the foot.

### Squirrel monkeys were not more likely to become infected as dose of ZIKV increased, duration of viremia was inversely related to peak titer, and transmission to mosquitoes increased with increasing viremia

For analyses of the magnitude and duration of viremia, we chose to exclude 4683, the animal euthanized at 13 dpi (**Supplemental Text S.6**), as we did not know whether pre-existing health problems may ultimately have led to euthanasia and may have impacted viral replication or immune parameters; however, we note below when inclusion of this animal altered determination of significance of comparisons. Contrary to Hypothesis 1, total dose delivered did not affect peak titer of ZIKV (DF = 1, F = 0.02, P = 0.88). In agreement with Hypothesis 1, peak virus titer was negatively associated with the duration of infection for ZIKV, albeit the range of duration was quite limited (ordinal logistic regression, DF = 1, chi-squared = 7.4, P = 0.007, R^2^ = 0.17). When these analyses were repeated with 4683 included, relationships were qualitatively similar, but the association between peak titer and duration of infection was no longer significant. All infected squirrel monkeys transmitted ZIKV on at least one day (**Table 2**) and likelihood of transmission increased with virus titer (**Fig. 4**, this analysis included 4683). Of mosquitoes that fed on dpi 3 and 4, one batch of mosquitoes died, but all other monkeys infected at least one mosquito that salivated detectable virus. The mean rate of virus dissemination to saliva from fed mosquitoes was 63% (± 12%) (**Table S.3**).

**Figure 4.**
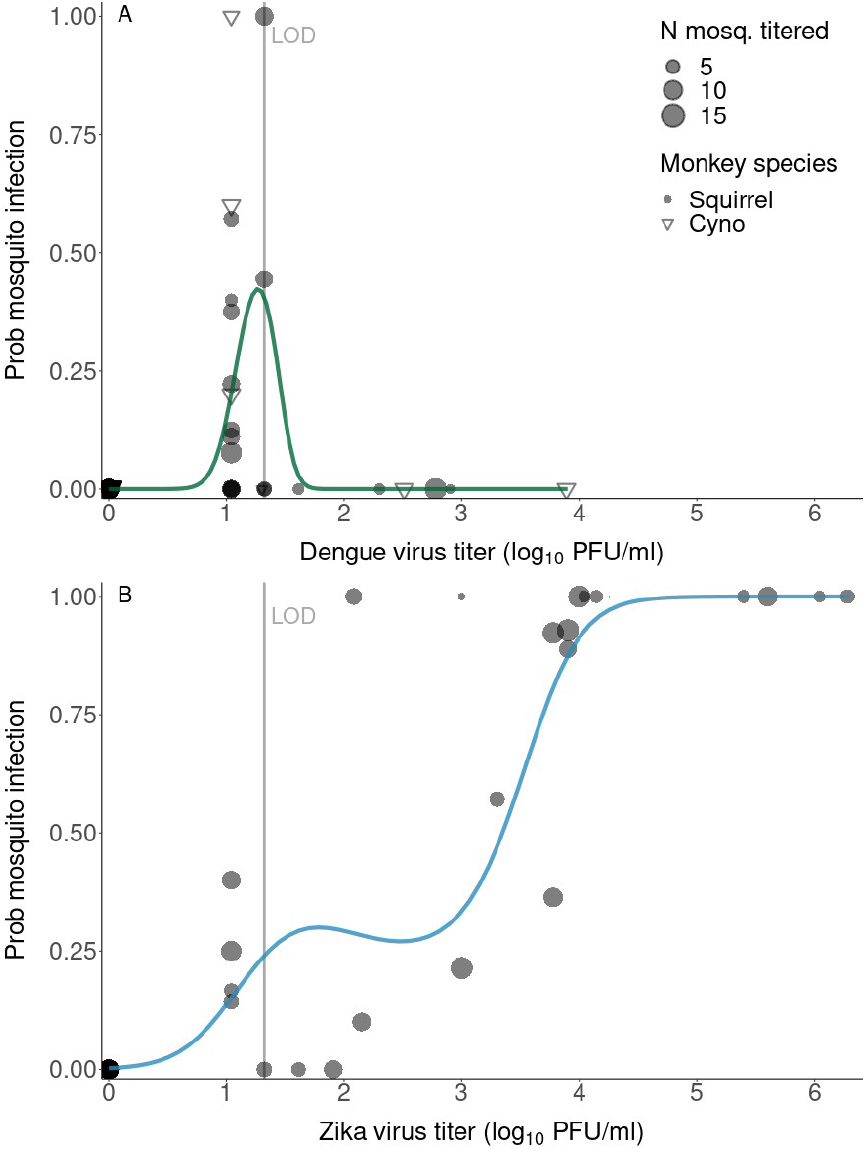
Relationship between virus titer in NHP serum and transmission of (A) sylvatic DENV-2 or (B) sylvatic ZIKV to *Ae. albopictus* (either body or leg infection). Curves fitted using a generalized additive model, allowing a variety of shapes, to unravel a possible pattern.

### In squirrel monkeys infected with ZIKV, higher early NK cells were associated with lower peak virus titer

Early % NK cells did not differ between ZIKV-infected and control monkeys (DF =11, t = -1.72, P = 0.11; **Fig. 3C**). Moreover, % NK cells in infected monkeys at early and later timepoints were significantly, positively correlated with each other (P = 0.005) but not with % NK cells prior to infection (P > 0.27 for both comparisons). Consistent with Hypothesis 2, higher early NK cells were associated with lower peak virus titer (DF = 1, F = 12.6, P = 0.009, adjusted R^2^ = 0.59 (**Fig. 3F**); this relationship was not significant when data from 4683 was included. It was not possible to analyze the association between early NK cells and neutralizing antibody titer, as approximately half of the PRNT80 values were <LOD.

### Comparison of sylvatic DENV-2 and sylvatic ZIKV dynamics in squirrel monkeys

Squirrel monkeys received significantly fewer bites from DENV-infected mosquitoes (3.4 (2.43 ; 4.76) bites) than ZIKV-infected mosquitoes (6.0 (4.66 ; 7.73) bites, P = 0.008, **Supplemental text S.4.1**) and the saliva titers of DENV-infected mosquitoes were significantly lower than those of ZIKV-infected mosquitoes (DENV 1.49 (1.38 ; 1.60), ZIKV 2.49 (2.30 ; 2.67) log_10_ pfu, P < 2e-16, **Supplemental text S.4.2**). Thus, squirrel monkeys received significantly lower doses of DENV than ZIKV (DENV 2.24 (1.98; 2.50) log_10_ pfu; ZIKV 3.47 (3.21 ; 3.74), P = 8.54e-11, **Supplemental text S.4.3**). Even though ZIKV peak titers were > 3 logs higher than DENV-2 titers (**Fig. 2, Tables 1 and 2**), PRNT80 values (with values <LOD set to 19) in ZIKV-infected monkeys were significantly lower (t-test, DF =18, t = - 4.65, P = 0.0002). Early % NK cells, in contrast, were similar between the two viruses (t-test, DF = 19, t = -0.101, P = 0.92) (**Fig. 3A, B, C**). A linear model comparing the relationship between virus dose and peak viremia between the two viruses revealed no interaction between virus and dose (DF = 1, F = 0.51, P = 0.49) and no effect of dose (DF = 1, F = 0.48, P = 0.50), but a highly significant effect of virus (DF = 1, F = 25.14, P = 0.00015), suggesting that the differences in dose delivered between the two viruses do not explain differences in peak titer. In a reduced dataset with similar range of doses for DENV and ZIKV ((2.3-3.3) log_10_ pfu; N = 5 DENV and 4 ZIKV), peak viremia was still significantly higher in ZIKV-than DENV-infected squirrel monkeys (DF = 1, F = 37.83, P = 4.7e-4, adjusted R^2^ = 0.82).

### Comparison of sylvatic DENV-2 and sylvatic ZIKV dynamics across both host species

Patterns of DENV-2 and ZIKV replication (**Fig. S.8**) were dramatically different. DENV-2 sustained low but prolonged viremia while ZIKV peaked quickly and was quickly cleared. Transmission also differed, as highlighted when fitting generalized additive models to each dataset (**Fig. 4, Supplemental text S.7**). The smooth term associated with virus titer was non-significant for DENV (p = 0.118) and significant for ZIKV (p <2e-16). The deviance explained by both models was 67.4% and.2% respectively.

### Relationship between viremia titer and transmission is similar between ZIKV in squirrel monkeys and DENV in humans

The best dose-response fit to characterize the relationship between viremia and transmission (mosquito body infection) of ZIKV in squirrel monkeys (the only host-virus pair with adequate levels of transmission for analysis) was obtained using a logistic equation (**Eq. S.1**) and a betabinomial likelihood (**Supplemental text S.8, Fig. S.9, Table S.7, S.8**). This revealed an inflection in rate of transmission at approximately 5 log_10_pfu/ml (**Fig. 5**). We compared this relationship to data generated by Nguyen et al. ^41^ in which viremia in infected humans and transmission to *Ae. aegypti* mosquitoes (measured as infection of mosquito bodies) was quantified (**Fig. 5**). For these data, the best fit was also a logistic equation using a betabinomial likelihood (**Supplemental text S.8, Fig. S.9**). We conducted a similar comparison to data on DENV transmission from infected humans to *Ae. aegypti* generated by Duong et al. ^42^ focusing on transmission measured as infection of mosquito legs (**Supplemental text S.8, Fig. S.9, S.10, Table S.7, S.8**). Interestingly, while the infectious dose 50 (ID_50_) for transmission to mosquito bodies was slightly higher for ZIKV than DENV, the ID_50_ for transmission to legs was substantially lower. Similarly, in our study, we found that a much higher proportion of ZIKV than DENV infections could be detected in mosquito legs than bodies (**Table S.6**).

**Figure 5.**
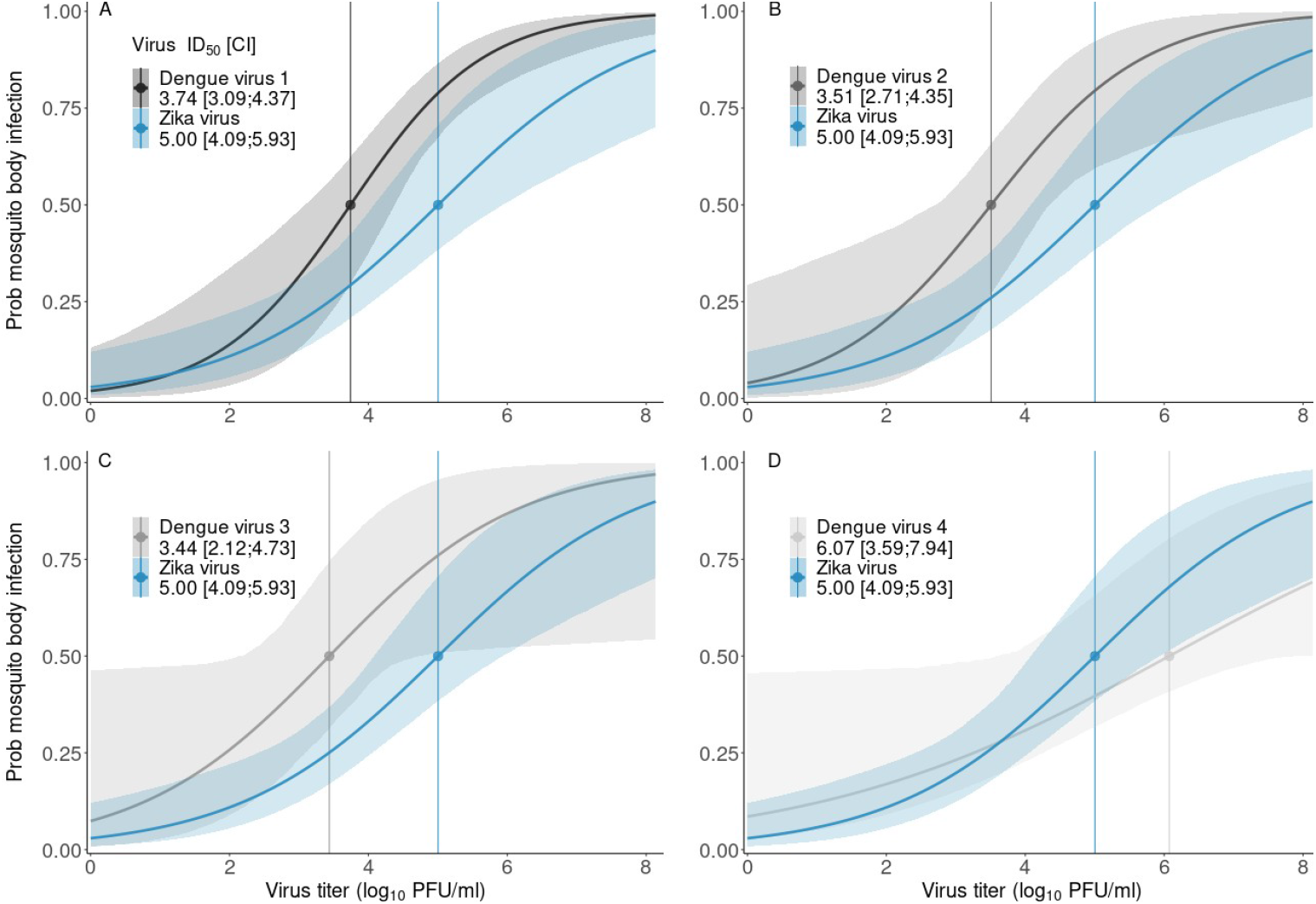
Relationship between virus titer in serum and transmission of ZIKV to bodies (sans legs) of *Ae. albopictus* (blue curves, same curve repeated in each panel) or transmission to abdomens of *Ae. aegypti* of (A) DENV serotype 1, (B) DENV serotype 2, (C) DENV serotype 3, (D) DENV serotype 4 (DENV data from Nguyen et al. 2013 ^41^). Infectious dose 50 (ID_50_), and confidence interval (CI), provided for each designated virus and mosquito species in log_10_ pfu/ml. Model selection with three competing functional forms, all with a sigmoidal shape.

## Discussion

This study tested predictions stemming from three hypotheses about replication-clearance trade-offs for arboviruses. First, we predicted that, if replication and clearance are locked into a trade-off, increasing doses of virus would result in higher virus peak titer, transmission would increase with virus titer, and higher peak titers would result in more rapid clearance of virus. Virus dose was estimated by summing the amount of virus in forced salivations from each mosquito that had fed upon a particular animal. While forced salivations likely underestimate the absolute amount of virus delivered ^43,44^, they should reflect the relative dose of virus.

Seroconversion revealed that sylvatic DENV-2 infected all nine cynomolgus macaques, a species known to become infected with the virus in nature, that received an infectious mosquito bite. However, only three monkeys produced viremia that could be detected directly from serum, and this viremia was only detectable on one or two days. Three macaques also transmitted virus to mosquitoes, but only one was a monkey from which viremia had been detected in serum, and viremia was not detected on the day of transmission. Similar patterns of muted but prolonged replication of DENV-2, and transmission from animals with undetectable/barely detectable viremia were observed in squirrel monkeys, a neotropical species for which no natural DENV infections have been documented. Higher virus doses increased like-lihood of detecting viremia in macaques but not squirrel monkeys; it was not possible to analyze peak titer in either species due to small number monkeys with quantifiable viremia.

In contrast to DENV-2, sylvatic ZIKV reached high titers in squirrel monkeys and mosquito transmission was positively related to virus titer. In infected mosquitoes, ZIKV disseminated efficiently to the saliva. Despite supporting high levels of viremia and transmission of ZIKV, squirrel monkeys raised neutralizing antibody titers that were significantly lower than those of DENV-2, and often below the LOD when a stringent cutoff (PRNT80) was used. Similarly, studies on Ross River virus (reviewed in ^45^) showed that experimentally infected bats produced no detectable viremia but nonetheless infected mosquitoes, while corellas (*Cacatoes correla*) became viremic but did not seroconvert. In combination, these findings highlight the complexity in using serological data to discern the role of species in virus transmission cycles ^46,47^.

A trade-off between magnitude and duration of infection was evident both within and between viruses. Within ZIKV infections, maximum virus titer was associated with shorter duration of infection. Comparing between viruses, ZIKV titers were significantly higher than DENV-2 titers in squirrel monkeys, but the monkeys infected with ZIKV were viremic for only three to seven days while the monkeys with detectable DENV-2 viremia were viremic over twelve to fourteen days. Interestingly, the effect of virus titer on transmission to mosquitoes was different for DENV and ZIKV, with DENV transmission occurring when viremia was low or just above the LOD, while ZIKV transmission was positively associated with titer. It seems that sylvatic DENV follows a “tortoise” strategy in squirrel monkeys, while sylvatic ZIKV follows a “hare” strategy ^29^.

Second, we predicted that, if NK cells play a role in shaping a replication-clearance trade-off, then higher mobilization of NK cells early in infection should suppress peak virus titer, which should in turn lower neutralizing antibody titers. For sylvatic DENV-2, peak titer could not be analyzed, but consistent with our prediction, as the percentage NK cells out of total PBMCs on early dpi increased, PRNT80 values measured at 28 dpi decreased. In squirrel monkeys infected with ZIKV the relationship with PRNT80 values could not be analyzed. Thus, there is substantial support that NK cells may play a direct role in shaping the replication of these viruses ^48^ or may be linked to another pathway that enacts this function. Intriguingly, we detected no consistent elevation in any of the cytokines measured concomitant with this shift in NK cells.

Third, we predicted that transmission from novel hosts would be low relative to native hosts, reflecting co-adaptation of viruses with their native hosts. Infection of both cynomolgus macaques and squirrel monkeys with sylvatic DENV via mosquito bite resulted in muted virus replication. Previous studies utilizing needle delivery of human-endemic DENV have reported peak virus titers about three orders of magnitude lower in cynomolgus macaques than neotropical primates ^28^. *Prima facie*, these latter data would suggest that transmission is actually more efficient from neotropical monkeys than Old World monkeys. However, our startling finding that sylvatic DENV transmission to mosquitoes occurred predominantly at undetectable or barely detectable viremia indicates a more complex relationship between viremia and transmission. Nonetheless, transmission of sylvatic DENV-2 from both macaques and squirrel monkeys was infrequent. This paucity of transmission could be the byproduct of a mismatch in the genetic interaction between sylvatic DENV-2 and *Ae. albopictus* Galveston ^49^. Alternatively, it could suggest that cynomolgus macaques are not actually a reservoir host for this virus. The paradigm that Asian NHPs are the reservoir of sylvatic DENV, rather than an amplifying host or a dead-end host ^50^, rests on scanty evidence, including serosurveys and isolation of sylvatic DENV from sentinel cynomolgus macaques ^34-36^. On the other hand, the lack of pathogenesis of sylvatic DENV-2 in macaques, the long duration of viremia, and indeed the low magnitude of viremia itself may be evidence that macaques are indeed a reservoir host ^51^. We will attempt to parameterize mathematical models with these data to assess whether or not cynomolgus macaques could be expected to sustain transmission of sylvatic DENV.

Similar to DENV, a Puerto Rican strain of ZIKV delivered by needle replicated to peak titers that were 2-3 orders of magnitude higher in neotropical marmosets and tamarins compared to rhesus and cynomolgus macaques ^52^. Cynomolgus macaques were not infected with ZIKV in the current study, but Dudley et al. ^30^ found that, compared to needle delivery, mosquito bite delivery of a Puerto Rican strain of ZIKV delayed the onset of viremia and delayed peak of viremia from day 3 to day 5 pi, but for both methods of delivery, the magnitude of virus replication was low (approximately 10^4^ pfu/ml at peak) and almost none of the *Ae. aegypti* fed upon these monkeys became infected. We reported similarly low transmission from cynomolgus macaques infected with a Cambodian strain of ZIKV via needle delivery and fed upon by *Ae. aegypti* ^53^. Thus, opposite our initial hypothesis, evidence to date suggests that neotropical primates support similar or higher levels of DENV and ZIKV transmission than their Old World counterparts. Moreover, similar to previous findings by Carrington et al. ^54^, we detected little difference in replication dynamics of DENV-2 and ZIKV after delivery by mosquito bite and studies in which the viruses were delivered by needle.

Finally, our data offer insights into two other central questions: why has DENV never spilled back into a sylvatic cycle in the Americas and what is the likelihood that ZIKV will do so? The rarity of transmission of sylvatic DENV-2 from squirrel monkeys suggests that the virus cannot gain a foothold in neotropical species, albeit testing more strains of mosquitoes and monkeys, as well as modeling, are still needed. In contrast, the viremia-transmission curves of sylvatic ZIKV in squirrel monkeys were similar to those derived from studies of DENV-1-4 in humans ^41,42^. ZIKV also reached higher titers than DENV in *Ae. albopictus* saliva after intrathoracic inoculation, consistent with results of a study by Chaves et al. ^55^ in which *Ae. aegypti* were infected with both viruses via membrane feeding. While additional modeling will be needed to draw inferences about the epidemic potential of *Ae. albopictus* for ZIKV in neotropical monkeys, as was done by Lequime et al. 2020 for a human population ^56^, our empirical data suggest that establishment of a neotropical, enzootic cycle is substantially more likely for ZIKV than DENV.

## Methods

### Viruses and cell lines

African green monkey kidney cells (Vero) and larval *Ae. albopictus* cells (C6/36) were purchased from the American Type Culture Collection (ATCC, Bethesda, MD, USA). Vero cells were maintained in Dulbecco’s Modified Eagle’s Medium (DMEM, ThermoFisher Scientific, Waltham, MA, USA) supplemented with 5% (v/v) heat-inactivated fetal bovine serum (FBS, Atlanta Biologicals, Flowery Branch, GA, USA), 1% (v/v) Penicillin–Streptomycin (P/S, ThermoFisher Scientific, Waltham, MA, USA; 100 U/mL and 100 μg/mL, respectively) in a humidified 37 °C incubator with 5% CO_2_. C6/36 cells were maintained in DMEM supplemented with 10% (v/v) heat-inactivated FBS, 10% (v/v) tryptose phosphate broth (TBP, Sigma-Aldrich, St. Louis, MO, USA), 1% (v/v) P/S in a humidified 28 °C incubator with 5% CO_2_.

The following viruses with the following passage histories were utilized to infect monkeys: sylvatic DENV-2 strain P8-1407 (suckling mouse (SM) 3 passages, C6/36 5 passages) and sylvatic ZIKV strain DakAR 41525 (AP-61 1 passage, C6/36 2 passages, Vero 6 passages). Pre-study blood samples were screened via PRNT assays against DENV-1 Hawaii, DENV-2 NGC, DENV-3 H87, DENV-4 Dominica, ZIKV PRVABC-59, and YFV 17D; all were negative (**Supplemental Table S.4**).

### Virus quantification from cell culture supernatants, NHP sera and mosquitoes

Virus stocks and NHP sera were titered in Vero cells while mosquito samples were titered in C6/36 cells. All NHP sera were titered starting at a 1:10 dilution and were also passaged once in Vero cells for 5-7 days as follows: generally, 500 μL of 1:10 diluted NHP serum in Vero cell media was used to infect one well of confluent Vero cells in 6-well plates, plates were gently rocked for two hours before adding media to a total volume of 3 mL and then plates were incubated for 4-6 days, cell supernatants were clarified by centrifugation, stabilized in 1X SPG (final concentration: 218 mM sucrose, 6 mM L-glutamic acid, 3.8 mM monobasic potassium phosphate, and 7.2 mM dibasic potassium phosphate, pH 7.2), and stored at - 80°C. For a relatively small number of serum samples with volumes <50 μL, dilutions of 1:20, 1:40 and 1:100 were used as necessary. Resulting cell supernatants were titered in Vero cells. Fifty μl of saliva samples were passaged once in one well of a 96-well plate of confluent C6/36 cells for 5-7 days, and the resulting cell supernatants were titered in C6/36 cells.

Viral quantification of cell-free viral stocks for infection and PRNT assays was conducted using standard infectious assays ^53^. Briefly, viral stocks underwent 10-fold serial dilutions using dilution media (DMEM, 2% FBS, and 1% P/S) in sterile 96-well tissue culture plates. Dilutions were added to 85-95% confluent monolayers of either Vero or C6/36 cells, as specified, in 12 or 24 well tissue culture plates. Dilutions were allowed to adsorb onto monolayers for 1 hour in a humidified 37 °C (Vero) or 28 °C (C6/36) incubator with 5% CO_2_ and were rocked every 15 minutes to prevent drying of the monolayers. Following adsorption, monolayers were overlayed using a solution of DMEM containing 3% FBS, 1% Pen–Strep, 1.25 μL/mL amphotericin B, and 0.8% weight/vol methylcellulose. The overlayed plates were incubated for 5 days in a humidified 37 °C incubator with 5% CO2; then, each well was washed twice with PBS and fixed for a minimum of 30 min in an ice-cold solution of methanol:acetone, 1:1, vol/vol. The fixative was removed and the plates were air dried. Following complete air drying, the plates were washed with PBS and then blocked with 3% FBS in PBS, followed by an overnight incubation with mouse hyperimmune serum against DENV-2 NGC or mouse hyperimmune serum against ZIKV strain MR-766 (1:2000 in blocking solution) or pan-flavivirus monoclonal antibody 4G2 (1:2000) at a 1:2000 dilution; mouse hyperimmune antibodies were obtained from the World Reference Center on Emerging Viruses and Arboviruses WRCEVA) at UTMB while 4G2 was obtained from VWR (Radnor, PA). The plates were then washed with PBS, followed by incubation with a goat anti-mouse secondary antibody conjugated to horseradish peroxidase (KPL, Gaithersburg, MD, USA) diluted 1:2000 in blocking solution. The plates were washed with PBS, after which an aminoethylcarbazole solution (Enzo Diagnostics, Farmingdale, NY, USA) prepared according to the manufacturer′s protocol was added, and the plates were incubated in the dark. Development was halted by washing in tap water, and the plates were allowed to air dry at room temperature before scoring. Virus quantification from monkey sera and mosquitoes was conducted using standard infectious assays, which are extremely similar to those described above save that titration was conducted in 24-well plates, plates were developed in 90% methanol at room temperature, incubated with primary antibodies for one hour, and developed using TrueBlue (SeraCare, Milford, MA, USA).

### Mosquito strains, maintenance, infection, and forced salivation

*Aedes albopictus* were field collected in Galveston Texas in the summer of 2018 and utilized to establish a continuous colony. All mosquitoes utilized over the course of these experiments were derived from this original colony. In Experiment 1, *Ae. albopictus* Galveston F12 were used, while in Experiment 2, *Ae. albopictus* Galveston F14 were used. All mosquitoes were maintained and/or utilized in Biosafety Level 2 facilities (Arthropod or Animal Containment) following standard procedures ^57^. After hatching, larvae were maintained in water in plastic containers, and then transferred after eclosion into 0.5L cardboard cups overlayed with mesh across the lids, and cartons were held in Tupperware containers within insect incubators set to 28 ± 1 °C with 80 ± 10% RH and a 16:8 light:dark cycle. Mosquitoes were provided *ad libitum* access 10% sucrose-soaked cotton rounds, save that one day prior to feeding on NHPs these were replaced with water-soaked rounds.

For mosquitoes used to infect monkeys, females at 4 days post-eclosion were IT inoculated with between 200 to 500 pfu of either DENV or ZIKV in a volume of 0.2 μL via a Nanoject II (Drummond Scientific, Broomall, PA, USA) and held in cartons as described. Ten days later, mosquitoes were cold-anesthetized and sorted into cartons based on treatment and individual NHP. Mosquitoes were then allowed to feed on monkeys as described below, engorged mosquitoes were separated and housed for 2-3 days. To harvest saliva, mosquitoes were cold anesthetized and immobilized on mineral oil, after which legs and wings were removed and the proboscis was inserted into sterile 10 μL pipette tips containing 10 μL of FBS for 30 minutes. After this, the saliva within the FBS was ejected into a microfuge tube containing 100 μL of DMEM containing 2% FBS, 1% P/S and 2.5 μg/mL of amphotericin B (hereafter referred to as mosquito collection media, MCM).

For mosquitoes used to detect transmission, following feeding, all mosquitoes were cold-anesthetized and engorged mosquitoes were separated and housed for 14 days. They were then cold anesthetized on ice and the legs of individual mosquitoes were removed; bodies and legs were placed in separate microfuge tubes containing sterilized steel ball bearings and 500 μL of MCM. In mosquitoes fed on squirrel monkeys at 3 and 4 days post-infection, saliva was also collected as described above.

### Non-human primate origins, maintenance, and monitoring

All procedures conducted on non-human primates were approved via UTMB Institutional Animal Care and Use Committee (IACUC) protocol 1912100. Mauritius origin adolescent *Macaca fascicularis* (between 2.4 to 5.4 years of age) weighing between 2.75-5.4 kg were purchased from Worldwide Primates, Inc (Miami, FL, USA). All animals were negative by serology for macacine alphaherpesvirus 1, simian immunodeficiency virus (SIV), simian type D retrovirus (SRV), simian T-lymphotropic virus 1 (STLV-1). Following a minimum 14-day quarantine, all animals were surgically implanted with DST micro-T temperature loggers (Star-Oddi, Garðabær, Iceland) set to record temperature every 15 minutes. Macaques were single housed in open metal caging that allowed for visual but not physical contact with other animals in the room. Standard primate chow was provided twice daily, with enrichment in the form of fruits and vegetables added once daily. All animals received a minimum of twice daily health checks.

*Saimiri boliviensis* boliviensis (Black capped squirrel monkeys) over an adolescent to adult age range (between 4 to 14 years of age) were purchased from the MD Anderson Center (Bastrop, TX, USA). Following a minimum 14-day quarantine, all animals were surgically implanted with DST micro-T temperature loggers (Star-Oddi, Garðabær, Iceland) set to record temperature every 15 minutes. Squirrel monkeys were pair housed in open metal caging that allowed for visual but not physical contact with animals in other cages. Standard primate chow was provided twice daily, with enrichment in the form of fruits and vegetables added once daily. All animals received a minimum of twice daily health checks.

### Non-human primate infection, blood sampling and mosquito feeding

All experimental procedures requiring any manipulation of the macaques were conducted following an overnight fast and under anesthesia, via intramuscular injection of 5-20 (cynomolgus macaques) or 10-15 (squirrel monkey) mg/kg ketamine. Once anesthetized, animals were removed from caging, and physical examination and weights were taken. To maintain core body temperature in squirrel monkeys following anesthesia, all procedures took place atop a preheated heating blanket. To infect monkeys, a 0.5L cardboard cup containing either 1, 10 or 15 (**Fig. 1**) sucrose starved *Ae. albopictus* Galveston previously injected with DENV, ZIKV or PBS was placed on the anesthetized animal’s ear and the mosquitoes were allowed to feed for 7-10 minutes. On subsequent time points following infection, cartons of 10 (Experiment 1) or 15 (Experiment 2) sucrose starved naïve *Aedes albopictus* were allowed to feed on the ear of the anesthetized animals.

Concurrent with mosquito feeding, blood was collected via venipuncture of the femoral vein. To measure cytokines, viremia and PRNTs, blood was aliquoted into serum separator tubes (SSTs, BD Biosciences) and centrifuged at 2000g for 5 minutes to pellet cells and clarified serum was aliquoted into fresh tubes. The others were retained for viremia and PRNT analysis. For quantification of NK cells, whole blood from venipuncture was added to lithium heparin microtainer tubes (BD Biosciences) and gently inverted.

### Natural killer cell quantification

Lithium heparin microtainer tubes containing 0.2 – 0.3 ml blood were taken into a biosafety cabinet and 200 μL of heparinized whole blood from each NHP was added to an individually labelled round-bottomed polystyrene tube (Corning/Falcon, Corning, NY, USA). Nonspecific antigen binding to Fc receptors was inhibited via 10 minutes of blocking with Human TruStain FcX (BioLegend, San Diego, CA, USA). Cells were subsequently stained with mouse anti-human CD20 (Alexa-700), mouse anti-human CD3 (Alexa-700), mouse anti-human CD14 (Alexa-700), mouse anti-human CD16 (PE) (all antibodies purchased from BD Biosciences), and Live/Dead Fixable Blue (Invitrogen, Waltham, MA, USA) in a volume of 100 μL for 1 hour in the dark with gentle agitation every 15 minutes. Subsequently, erythrocytes were lysed with BD FACS Lysing Solution (BD Biosciences) diluted in molecular biology grade water (Corning, Corning, NY, USA) for 12 minutes followed by immediate centrifugation at 500g for 5 minutes at room temperature. Supernatant was immediately aspirated and the cellular pellet was reconstituted in FACS Buffer (0.2% bovine serum albumin, 2mM EDTA in phosphate buffered saline). Samples were centrifuged at 350g for 5 minutes at room temperature. FACS buffer was removed and pellets were fixed in freshly prepared 2% methanol free formaldehyde (diluted from 16% Ultra Pure EM grade formaldehyde (Polysciences, Warrington, PA, USA) for a minimum of 1 hour prior to analysis. Analyses were conducted using a BD LSRFortessa Cell Analyzer (BD, Franklin Lakes, NJ, USA), and 10^5^ total cells were analyzed. Natural killer cells were defined as the live cell population that were CD16^+^ and CD3^-^CD14^-^ CD20^-^. Data analysis was conducted utilizing FlowJo software (FlowJo, Ashland, OR, USA).

### Cytokine quantification

Clarified serum from each animal was subjected to the Cytokine 29-Plex Monkey Panel (ThermoFisher Scientific, Waltham, MA, USA). Each sample was run in duplicate according to manufacturer instructions and average values were ascertained via the standard curve generated for each cytokine per run. Values that were outside the dynamic range of the assay were treated as being below LOD.

### Plaque reduction neutralization titers

PRNTs were performed in Vero cells in a 12-well tissue culture plate format as previously described ^58^. Briefly, pre-exposure serum and 28-day post-exposure serum samples were heat inactivated in a 56°C water bath for 60 minutes to inactivate complement. Serum samples subsequently underwent 2-fold serial dilution series in DMEM with 2% FBS and 1% P/S (1:10 to 1:320). An equal volume (75 μl) of ZIKV or DENV at a titer of 800 focus forming units (FFU) per milliliter was added to each serum dilution (dilution series range 1:20-1:640). Serum/virus mixtures were incubated for one hour at 37 °C, at which point 100 μL of mixture was added to a nearly confluent (85-95%) monolayer of Vero cells and allowed to adsorb for one hour at 37 °C. Each well was then overlayed DMEM containing 3% FBS, 1% Pen–Strep, 1.25 μL/mL amphotericin B, and 0.8% weight/vol methylcellulose and incubated 37 °C for 5 days. Fixation and immunostaining procedures took place as described above. PRNT titers were represented as the reciprocal of the highest dilution of serum that inhibited a specified percentage foci when compared to a virus only control.

### Statistical analysis and model fitting

To test *a priori* predictions within and between each species-virus combination, we used standard parametric tests or generalized linear models, as appropriate. *A priori* power analyses were conducted to assess our ability to detect changed in peak titer and duration of titer between low and high dose infections in macaques, resulting in a sample size of 10 in the infected group. As one macaque was transferred to the control group, and as analyses were primarily focused on NK cells and neutralizing antibody titers due to the paucity of viremia, we conducted a post-hoc test of power, and found that, with the effect sizes and standard deviations reported, we had approximately 75% power to detect a significant difference in comparisons among 2 or 3 groups and lower power of approximately 20% for linear regressions. Sample sizes were smaller for infected squirrel monkeys (N = 8 per virus) and power was concomitantly lower. For models testing interactions, if the interactions term was non-significant with an effect size η^2^ < 0.01, we relied on a type II anova for the significance of main effects, following recommendations by Smith and Cribbie ^59^.

Differences in cytokine concentration (log_10_ pg/μl) between sexes at baseline were assessed (**Supplemental Text S.1**). A possible long-term disruption of the cytokine equilibrium was assessed in infected individuals by comparing cytokine concentrations (log_10_ pg/μl) at baseline and at day 28 (**Supplemental Text S.2**). Tests were conducted with values below the LOD excluded or included (in that case, fixed at LOD), and compared. To account for multiple testing, we applied a *P* value threshold of 0.001 before considering a result further.

To further explore the drivers of cytokine concentrations, we used mixed effect models (**Supplemental text S3, S5** ^60^). We assessed differences in concentration between infected and control NHPs, as well as differences between viruses or NHP species. We used R packages glmmTMB ^61^ and gam ^62^ to run the models. All models used the log_10_ cytokine concentration as the response variable, but varied in their random effect (intercepts) structure (**Supplemental Text S.3.1, S.5**). All concentrations below LOD were excluded from analysis initially, for all models. This substantially improved the residuals of the models, which were checked using the R package DHARMA ^63^. To account for multiple testing, we applied a *P* value threshold of 0.001 before considering a result further. Results considered significant after this *P* value correction were compared to corresponding models including measures below LOD (in that case, fixed at LOD).

As previous work showed that cytokines such as IL-12, IL-15, and TNFα activate NK cells during flavivirus infection ^48,64^, we tested possible associations between the concentration of these cytokines (log_10_ pg/μl) and the percentage of NK cells, in infected and control NHPs, in the first week post-infection (**Supplemental text S.3.2**). Model selection was performed to discriminate between linear and generalized additive models (a more flexible fitting approach), as well as different random effect structures (**Supplemental text S.3.2**).

The number of infectious bites initially received per NHP, as well as the viral titers contained in biting mosquitoes’ saliva, are the components driving the initial dose delivered to NHPs, which can then drive their viral dynamics and immune response. Differences between experiments regarding those factors were assessed using generalized linear models (**Supplemental Text S.5**).

To quantify a possible relationship between host infectious titer (log_10_ pfu/ml) and probability to infect mosquitoes, we first used a generalized additive model. The probability of mosquito infection was broadly defined, measured by a positive mosquito body or leg. We used a binomial error distribution and constrained the number of knots to 6 to avoid overfitting. Separate relationships were fitted for dengue and Zika viruses, and transmission from both NHP species was considered for dengue.

We then refined the fitting of dose-response relationships between ZIKV infectious titers (log_10_ pfu/ml) of squirrel monkeys and the probability to infect mosquitoes. Separate relationships were fitted depending on whether mosquito infection was measured by the presence of virus in their abdomens or their legs (**Supplemental Text S.8**). In order to compare these ZIKV dose-response relationships to existing results on dengue virus transmission from humans to *Aedes aegypti*, we retrieved data from Nguyen et al. 2013 (for abdomen infections) ^41^ and Duong et al. ^42^ (for leg infections), through the supplementary material of ten Bosch et al. 2018 ^65^. As those data were in units RNA copies/ml, we applied a conversion factor, specific to each dengue serotype from Blaney et al. ^66^ (**Supplemental Text S.8**), to transform it into pfu/ml, and applied the same fitting procedure.

All analyses were performed in R and all scripts are available at (link to be placed here upon acceptance).

## Acknowledgements

We thank the members of the support staff at UTMB and NMSU, who worked diligently in the face of COVID-19 restrictions during the pandemic. This study was funded by NIH grant R01AI145918. Squirrel monkeys acquired from the UT MD Anderson Cancer Center, Michael E. Keeling Center for Comparative Medicine and Research were supported by NIH grant 5P40OD010938-36 prior to sale to UTMB.

## References

1 Swei, A., Couper, L. I., Coffey, L. L., Kapan, D. & Bennett, S. Patterns, Drivers, and Challenges of Vector-Borne Disease Emergence. Vector Borne Zoonotic Dis 20, 159–170 (2020).

2 Guth, S., Hanley, K. A., Althouse, B. M. & Boots, M. Ecological processes underlying the emergence of novel enzootic cycles: Arboviruses in the neotropics as a case study. PLoS Negl Trop Dis 14, e0008338 (2020).

3 Barrett, A. D. T. The reemergence of yellow fever. Science 361, 847–848 (2018).

4 Bryant, J. E., Holmes, E. C. & Barrett, A. D. Out of Africa: a molecular perspective on the introduction of yellow fever virus into the Americas. PLoS Pathog 3, e75 (2007).

5 Hanley, K. A. et al. Fever versus fever: the role of host and vector susceptibility and interspecific competition in shaping the current and future distributions of the sylvatic cycles of dengue virus and yellow fever virus. Infect Genet Evol 19, 292–311 (2013).

6 Aliota, M. T. et al. Zika in the Americas, year 2: What have we learned What gaps remain? A report from the Global Virus Network. Antiviral Res 144, 223–246 (2017).

7 Althouse, B. M. et al. Potential for Zika Virus to establish a sylvatic transmission cycle in the Americas. PLoS Negl Trop Dis 10, e0005055 (2016).

8 Vasilakis, N. & Weaver, S. C. Flavivirus transmission focusing on Zika. Curr Opin Virol 22, 30–35 (2017).

9 Favoretto, S. R. et al. Zika virus in peridomestic neotropical primates, Northeast Brazil. EcoHealth 16, 61–69 (2019).

10 Terzian, A. C. B. et al. Evidence of natural Zika virus infection in neotropical non-human primates in Brazil. Sci Rep 8, 16034 (2018).

11 Cardosa, J. et al. Dengue virus serotype 2 from a sylvatic lineage isolated from a patient with dengue hemorrhagic fever. PLoS Negl Trop Dis 3, e423 (2009).

12 Diallo, D. et al. Concurrent amplification of Zika, chikungunya, and yellow fever virus in a sylvatic focus of arboviruses in Southeastern Senegal, 2015. BMC Microbiol 20, 181 (2020).

13 Franco, L. et al. First report of sylvatic DENV-2-associated dengue hemorrhagic fever in West Africa. PLoS Negl Trop Dis 5, e1251 (2011).

14 Gubler, D. J., Vasilakis, N. & Musso, D. History and Emergence of Zika Virus. J Infect Dis 216, S860–S867 (2017).

15 Pyke, A. T. et al. Complete Genome Sequence of a Highly Divergent Dengue Virus Type 2 Strain, Imported into Australia from Sabah, Malaysia. Genome Announc 5 (2017).

16 Pyke, A. T. et al. Highly divergent dengue virus type 1 genotype sets a new distance record. Sci Rep 6, 22356 (2016).

17 Teoh, B. T., Sam, S. S., Abd-Jamil, J. & AbuBakar, S. Isolation of ancestral sylvatic dengue virus type 1, Malaysia. Emerg Infect Dis 16, 1783–1785 (2010).

18 Childs, L. M. et al. Linked within-host and between-host models and data for infectious diseases: a systematic review. PeerJ 7, e7057 (2019).

19 Visher, E. et al. The three Ts of virulence evolution during zoonotic emergence. Proc Biol Sci 288, 20210900 (2021).

20 Bull, J. J. & Lauring, A. S. Theory and empiricism in virulence evolution. PLoS Pathog 10, e1004387 (2014).

21 Frank, S. A. Models of parasite virulence. Q Rev Biol 71, 37–78 (1996).

22 Hanley, K. A. et al. Infection dynamics of sylvatic dengue virus in a natural primate host, the African Green Monkey. Am J Trop Med Hyg 91, 672–676 (2014).

23 Koide, F. et al. Development of a Zika Virus Infection Model in Cynomolgus Macaques. Front Microbiol 7, 2028 (2016).

24 Rayner, J. O. et al. Comparative Pathogenesis of Asian and African-Lineage Zika Virus in Indian Rhesus Macaque’s and Development of a Non-Human Primate Model Suitable for the Evaluation of New Drugs and Vaccines. Viruses 10 (2018).

25 Crooks, C. M. et al. African-Lineage Zika Virus Replication Dynamics and Maternal-Fetal Interface Infection in Pregnant Rhesus Macaques. J Virol 95, e0222020 (2021).

26 Raasch, L. E. et al. Fetal loss in pregnant rhesus macaques infected with high-dose African-lineage Zika virus. PLoS Negl Trop Dis 16, e0010623 (2022).

27 Alizon, S. & van Baalen, M. Transmission-virulence trade-offs in vector-borne diseases. Theor Popul Biol 74, 6–15 (2008).

28 Althouse, B. M. et al. Viral kinetics of primary dengue virus infection in non-human primates: a systematic review and individual pooled analysis. Virology 452-453, 237–246 (2014).

29 Althouse, B. M. & Hanley, K. A. The tortoise or the hare Impacts of within-host dynamics on transmission success of arthropod-borne viruses. Philos Trans R Soc Lond B Biol Sci 370 (2015).

30 Dudley, D. M. et al. Infection via mosquito bite alters Zika virus tissue tropism and replication kinetics in rhesus macaques. Nat Commun 8, 2096 (2017).

31 Le Coupanec, A. et al. Aedes mosquito saliva modulates Rift Valley fever virus pathogenicity. PLoS Negl Trop Dis 7, e2237 (2013).

32 Schneider, B. S. et al. Aedes aegypti saliva alters leukocyte recruitment and cytokine signaling by antigen-presenting cells during West Nile virus infection. PLoS One 5, e11704 (2010).

33 Ben-Shachar, R. & Koelle, K. Transmission-clearance trade-offs indicate that dengue virulence evolution depends on epidemiological context. Nat Commun 9, 2355 (2018).

34 Rudnick, A. Studies of the ecology of dengue in Malaysia: a preliminary report. J Med Entomol 2, 203–208 (1965).

35 Rudnick, A. Ecology of dengue virus. Asian J. Infect. Dis. 2, 156–160 (1978).

36 Rudnick, A. & Lim, T. W. Dengue virus ecology in Malaysia. Inst. Med. Res. Malays. Bull. 23, 51–152 (1986).

37 Pereira-Dos-Santos, T., Roiz, D., Lourenco-de-Oliveira, R. & Paupy, C. A Systematic Review: Is Aedes albopictus an Efficient Bridge Vector for Zoonotic Arboviruses? Pathogens 9 (2020).

38 Grard, G. et al. Zika virus in Gabon (Central Africa)--2007: a new threat from Aedes albopictus? PLoS Negl Trop Dis 8, e2681 (2014).

39 Laffins, M. M., Mellal, N., Almlie, C. L. & Regalia, D. E. Evaluation of Infrared Thermometry in Cynomolgus Macaques (Macaca fascicularis). J Am Assoc Lab Anim Sci 56, 84–89 (2017).

40 J, F., T, H. & B, B. The laboratory nonhuman primate. (CRC Press 2002).

41 Nguyen, M. N. et al. Host and viral features of human dengue cases shape the population of infected and infectious Aedes aegypti mosquitoes. Proc Natl Acad Sci U S A 110, 9072–9077 (2013).

42 Duong, V. et al. Asymptomatic humans transmit dengue virus to mosquitoes. Proc Natl Acad Sci U S A 112, 14688–14693 (2015).

43 Gloria-Soria, A., Brackney, D. E. & Armstrong, P. M. Saliva collection via capillary method may underestimate arboviral transmission by mosquitoes. Parasit Vectors 15, 103 (2022).

44 Miller, M. R. et al. Characterizing and Quantifying Arbovirus Transmission by Aedes aegypti Using Forced Salivation and Analysis of Bloodmeals. Insects 12 (2021).

45 Skinner, E. B. et al. Species Traits and Hotspots Associated with Ross River Virus Infection in Nonhuman Vertebrates in South East Queensland. Vector Borne Zoonotic Dis 21, 50–58 (2021).

46 Gilbert, A. T. et al. Deciphering serology to understand the ecology of infectious diseases in wildlife. Ecohealth 10, 298–313 (2013).

47 Viana, M. et al. Assembling evidence for identifying reservoirs of infection. Trends Ecol Evol 29, 270–279 (2014).

48 Bjorkstrom, N. K., Strunz, B. & Ljunggren, H. G. Natural killer cells in antiviral immunity. Nat Rev Immunol 22, 112–123 (2022).

49 Lambrechts, L. et al. Genetic specificity and potential for local adaptation between dengue viruses and mosquito vectors. BMC Evol Biol 9, 160 (2009).

