## Supplementary Information for "Immunologically mediated trade-offs shaping transmission of sylvatic dengue and Zika viruses in native and novel non-human primate hosts"

Note : Tables S1 to S6 are provided as separate files

#### Temperature and weight of non-human primates over the course of the experiment

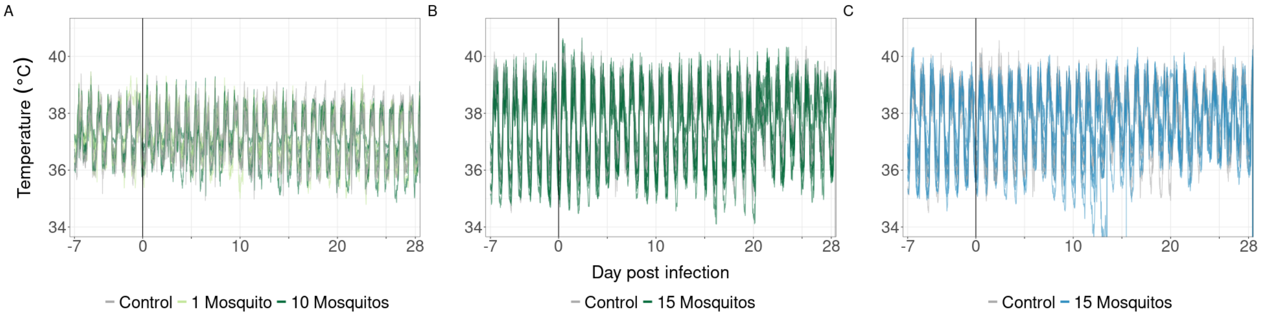


Figure S.1: Changes in temperature in (A) cynomolgus macaques infected with DENV (green lines) or control (grey lines), (B) squirrel monkeys infected with DENV (green lines) or control (grey lines), and (C) squirrel monkeys infected with ZIKV (blue lines) or control (grey lines). Note that the same set of four control animals is shown in panels B and C and that the two animals in panel C that drop below 34°C were the two animals that were euthanized over the course of the experiment. Normal temperature ranges are [36;39.5]°C for cynomolgus macaques[1,2] and [33.3;41]°C for squirrel monkeys[3,4].

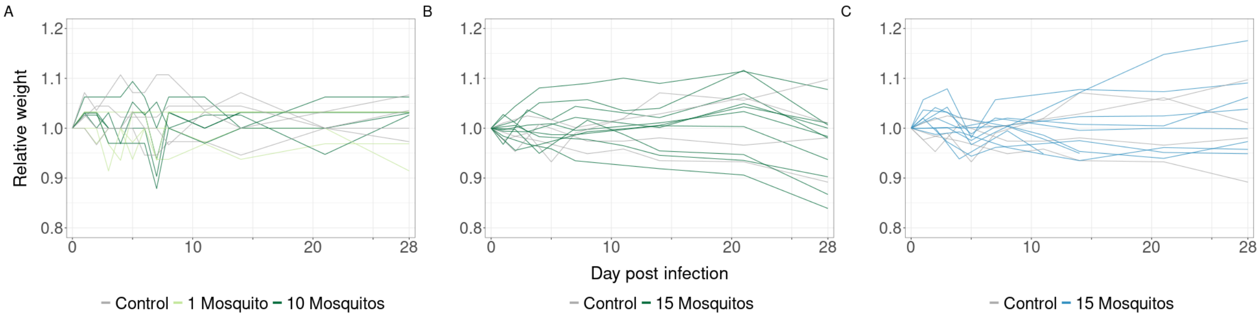


Figure S.2: Changes in body weight in (A) cynomolgus macaques infected with DENV (green lines) or control (grey lines), (B) squirrel monkeys infected with DENV (green lines) or control (grey lines), and (C) squirrel monkeys infected with ZIKV (blue lines) or control (grey lines). Note that the same set of four control animals is shown in panels B and C.

#### S.1 Cytokines - Sex differences at baseline

To test possible sex differences at baseline in $\log_{10}$ cytokine concentrations, we performed Wilcoxon tests, preceded by Levene's tests to check homogeneity of variance. This was done for each cytokine and each NHP species separately. Tests were conducted with values below the limit of detection (LOD) excluded or included (in that case, fixed at LOD), and compared. To account for multiple testing, we set alpha (the threshold for significance) to 0.001.
Detailed results can be found in Table S.5.1. No significant differences between sexes at baseline were found.

#### S.2 Cytokines - Long-term disruption of the cytokine equilibrium

Long-term disruption of the cytokine equilibrium was assessed in infected individuals by comparing cytokine concentrations ($\log_{10}$) at baseline and at day 28, for each arm of the experiments (DENV-infected macaques, DENV-infected squirrel monkeys, ZIKV-infected squirrel monkeys) and each cytokine separately, using a paired Wilcoxon test. When a difference was detected for a given cytokine, it was qualitatively compared to the change observed in control individuals of the same species. Tests were conducted with values below the limit of detection (LOD) excluded or included (in that case, fixed at LOD), and results were compared. To account for multiple testing, we set alpha (the threshold for significance) to 0.001.
Differences in cytokine concentration between baseline (day -7) and day 28 in infected individuals were detected in cynomolgus macaques for I-TAC, IL-12, and IL-RA (Table S.5.2), but $P$ values did not meet our threshold for multiple testing. Nevertheless, data showed that similar differences could be observed in control individuals, excluding a possible long-term effect of infection.

#### S.3 Cytokines - Effect of infection

##### S.3.1 Differences in cytokine dynamics between infected and control NHPs

We assessed possible differences in concentration between infected and control NHP, in each arm of the experiments (DENV in macaques, DENV in squirrel monkeys, ZIKV in squirrel monkeys), and for each cytokine separately. To do so, we fitted linear mixed effect models using maximum likelihood, with the $\log_{10}$ cytokine concentration as the response variable, the group (infected or control) as the fixed effect, and day and individual as random effects. All concentrations below the limit of detection (LOD) were excluded from analysis initially, for all models. This substantially improved the residuals of the models, which were checked using the R package DHARMA. To account for multiple testing, we set alpha (the threshold for significance) to 0.001. Results considered significant after this $P$ value correction were compared to corresponding models including measures below LOD (in that case, fixed at LOD). Cytokine concentrations measured before infection and at day 28 post infection were excluded from analyses.
The concentration of RANTES was significantly higher in controls than in ZIKV-infected squirrel monkeys (Figure S.7 *, Table S.5.3). For the reference model (excluding <LOD measures), we estimate an increase of 0.27 [0.14 ; 0.41] $\log_{10}$ pg/µl, $P$ value 8.82e-5. For the model including <LOD measures, we estimate an increase of 0.32 [0.20 ; 0.44] $\log_{10}$ pg/µl, $P$ value 1.6e-7.

##### S.3.2 Effect of specific cytokines on NK cell activation

As previous work showed that cytokines such as IL-12, IL-15, and TNFα activate NK cells during flavivirus infection (Marquardt et al. 2015[5], Björkström et al. 2022[8]), we aimed to test possible associations between the concentration of these cytokines ($\log_{10}$ pg/µl) and the percentage of NK cells, in infected and control NHPs, in the first week post-infection. In cynomolgus macaques, the number of measures below LOD for IL-15 (20/28 in controls, 60/63 in DENV-infected) and for TNFα (23/28 in controls, 50/63 in DENV-infected) was too important to fit desired models. In squirrel monkeys, which had less timepoints with both cytokines and percentage NK cells measured, the number of measures below LOD for IL-15 (8/8 in controls, 14/20 in ZIKV-infected, 14/20 in DENV-infected), for TNFα (8/8 in controls, 17/20 in ZIKV-infected, 19/20 in DENV-infected), and for IL-12 (8/8 in controls, 13/20 in ZIKV-infected, 13/20 in DENV-infected) was too important to fit desired models. We therefore focused on IL-12 in cynomolgus macaques.

*This figure is placed later in the document to respect sequential citations in the manuscript. Sorry for the inconvenience

For each population (control and DENV-infected), we selected between linear and generalized additive models (gam), with or without random effects on day and/or monkey ID (random intercept only or random intercept and slope). In linear models, random intercepts and slopes were uncorrelated, to be consistent with what could be fitted in a gam. In gam models, the number of knots was constrained to 4 for the effect of cytokine concentration, and 2 for random effects, to avoid overfitting. The models were fitted using maximum likelihood, and the selection was based on likelihood ratio tests and corrected Akaike Information Criterion (AICc) comparison.
In both populations, the selected model was a gam including random slopes and intercepts per monkey ID, and random intercepts per day. There was no significant association between IL-12 concentration and % NK cells (P = 0.92 in controls, P = 0.075 in DENV-infected, Figure S.3). We note substantial uncertainty around the fits and strong heterogeneity between individuals (Figure S.3).

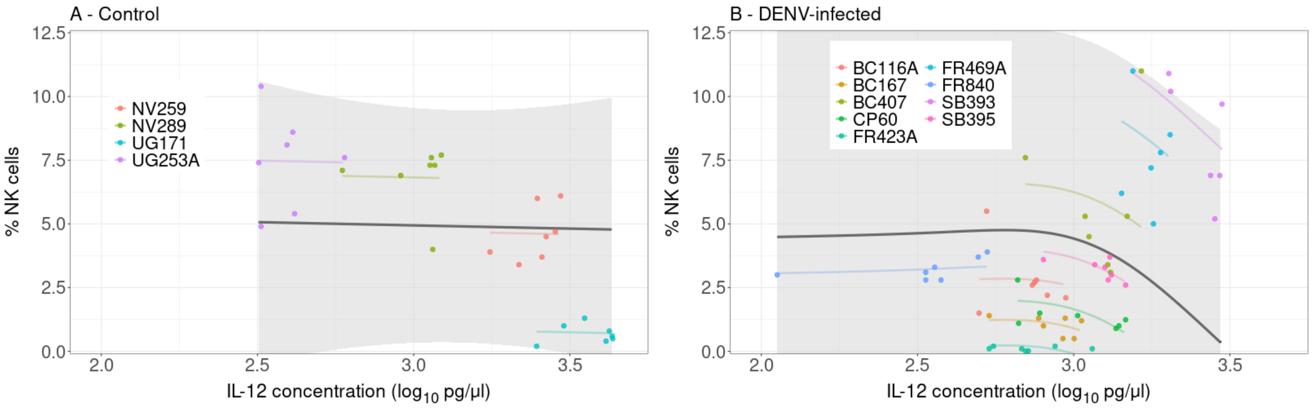


Figure S.3: Effect of IL-12 concentration on % NK cells in control (A) and DENV-infected (B) cynomolgus macaques. Results from generalized additive models, estimating an intercept and a slope per monkey. Grey lines and shading show the population trends with uncertainty. Points show data, with a color per monkey, and the associated fits with a line of the same color. Note that similar colors do not refer to the same individuals in panels A and B.

Even though this is not highlighted in the selected model, we note that there seem to be contrasting patterns of association between IL-12 concentration and % NK cells in control *vs* DENV-infected cynomolgus macaques, when looking at daily data (negative or bell-shaped in controls, positive in DENV-infected, Figure S.4). These patterns are not present within-individual, and do not seem driven by detectable viremia or transmission to mosquitoes (Figure S.4B). There might be a hidden process driving these associations, that should be investigated further.

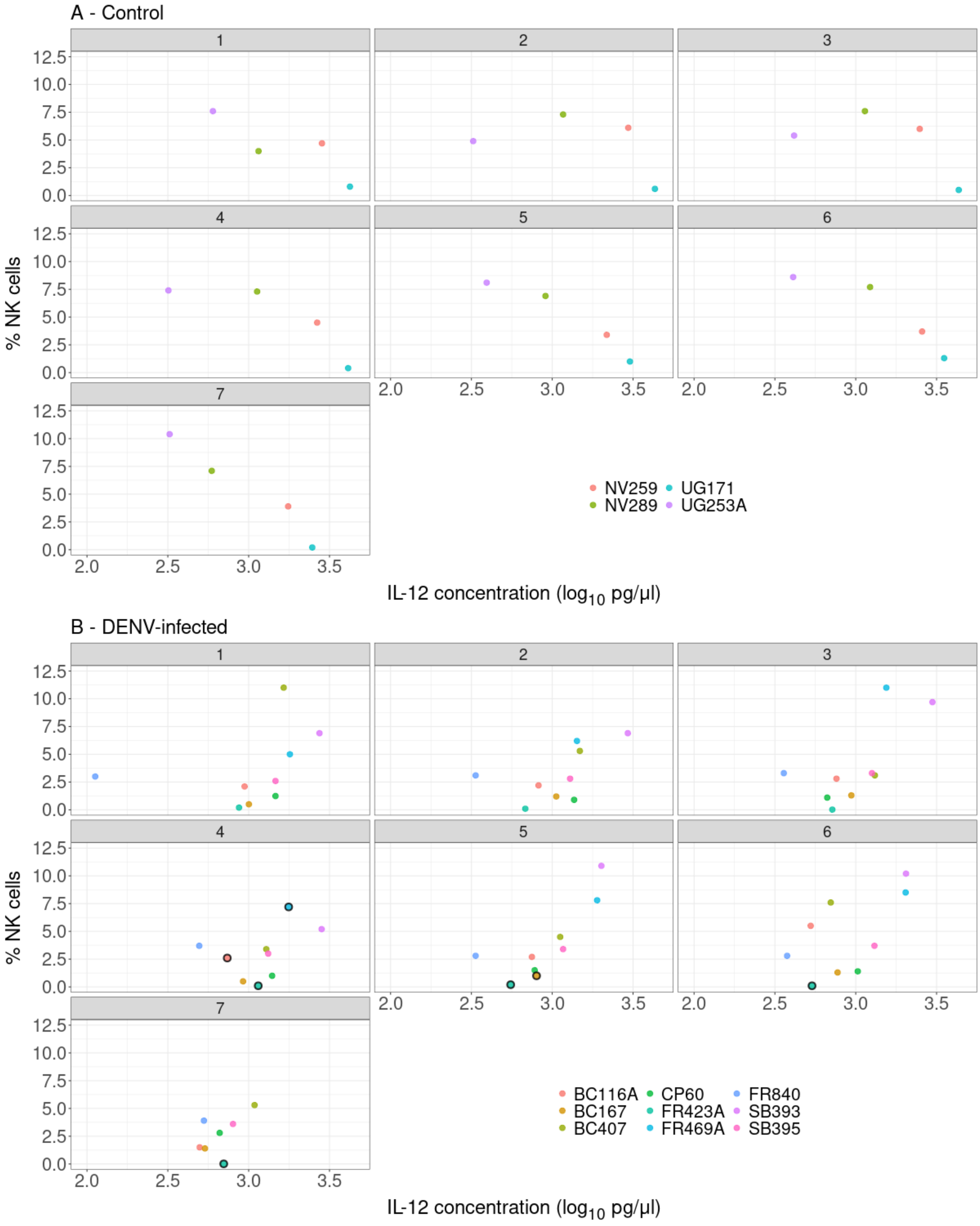


Figure S.4: Data on IL-12 concentration and % NK cells in control (A) and DENV-infected (B) cynomolgus macaques, per monkey (color) and timepoints (sub-panels). Black circled points indicate monkey and timepoints for which there was detectable viremia or transmission to mosquitoes. Note that similar colors do not refer to the same individuals in panels A and B.

#### S.4 Differences in dose delivered to NHPs on day 0

The number of infectious bites initially received per NHP, as well as the viral titers contained in biting mosquitoes' saliva, are the components driving the initial dose delivered to NHPs, which can then drive their viral dynamics and immune response. Differences between experiments regarding those factors were assessed using generalized linear models (Sections S.4.1-S.4.3).
For models using saliva titers of IT-injected *Ae. albopictus* (either individual titers or their sum per NHP), we note that assigning a random value between 0 and 39 PFU/collection for samples positive only after passage in C6/36 cells resulted in better model residuals than a fixed value of 20 (half the limit of detection). But the coefficients estimates and *P* values were not strongly impacted, so we kept the latter approach for simplicity.

##### S.4.1 Number of infectious bites

The proportion of positive bites was computed based on mosquitoes that fed and survived to titer. We applied this proportion to the number of mosquitoes that fed, and rounded to the nearest integer, to estimate the true number of positive bites each NHP got, which was our response variable. We then used a generalized linear model, with a Poisson error distribution and a log link function, to detect differences between experiments (defined by NHP species and virus, dengue virus and squirrel monkeys used as reference). We excluded mosquitoes belonging to the low exposure group feeding on cynomolgus macaques. The *P* value threshold was 0.05.
Squirrel monkeys initially received a similar number of DENV-infectious bites than cynomolgus macaques from the high exposure group (squirrel 3.4 [2.43 ; 4.76] infectious bites, cyno 4.2 [2.74 ; 6.44], *P* value 0.45), but significantly less than ZIKV-infectious bites received by squirrel monkeys (6 [4.66 ; 7.73], *P* value 0.0082).

```
 Family: poisson  ( log )
Formula:          pos_corr ~ group
Data: df_high

     AIC      BIC   logLik deviance df.resid
   107.4    111.1    -50.7    101.4       22


Conditional model:
                         Estimate Std. Error z value Pr(>|z|)
(Intercept)                1.2238     0.1715   7.136 9.62e-13 ***
groupCyno.Dengue virus     0.2113     0.2775   0.761  0.44645
groupSquirrel.Zika virus   0.5680     0.2147   2.646  0.00815 **
---
Signif. codes:  0 ‘***’ 0.001 ‘**’ 0.01 ‘*’ 0.05 ‘.’ 0.1 ‘ ’ 1
```

##### S.4.2 Saliva titer per mosquito

We used a linear model to detect differences in $\log_{10}$ saliva titers of mosquitos (response variable) depending on the virus they were infected with, (fixed effect). DENV-infected mosquitoes (reference) were grouped together irrespective of which host species they fed on. This was run on positive saliva titers only. For those positive only after passage in C6/36 cells, a value of 20 PFU was assigned. The *P* value threshold was 0.05.
After residual check, we selected a model correcting heteroskedasticity (by adding a dispersion model), even though issues remained in the residuals. The effect estimate and its significance were similar to the approach using random values for measures below the limit of detection, which showed no residuals' issues.
The saliva titers of ZIKV-infected *Ae. albopictus* were significantly higher than those of DENV-infected *Ae. albopictus* (ZIKV 2.49 [2.30 ; 2.67] $\log_{10}$ PFU, DENV 1.49 [1.38 ; 1.60], *P* value <2e-16).

```
 Family: gaussian  ( identity )
Formula:          log_titer ~ virus
Dispersion:                 ~virus
Data: df_fixLOD
```

```
     AIC      BIC   logLik deviance df.resid
     196      207      -94      188      110


Conditional model:
                Estimate Std. Error z value Pr(>|z|)
(Intercept)       1.4869     0.0558  26.648   <2e-16 ***
virusZika virus   0.9982     0.1101   9.069   <2e-16 ***
---
Signif. codes:  0 ‘***’ 0.001 ‘**’ 0.01 ‘*’ 0.05 ‘.’ 0.1 ‘ ’ 1

Dispersion model:
                Estimate Std. Error z value Pr(>|z|)
(Intercept)      -1.7467     0.1890  -9.242  < 2e-16 ***
virusZika virus   1.0968     0.2649   4.140 3.48e-05 ***
---
Signif. codes:  0 ‘***’ 0.001 ‘**’ 0.01 ‘*’ 0.05 ‘.’ 0.1 ‘ ’ 1
```

##### S.4.3 Dose delivered to NHPs

We summed $\log_{10}$ saliva titers of mosquitoes biting a same NHP to estimate the dose delivered, which was our response variable. We then used a linear model to detect differences between experiments, with the distinction between exposure groups in cynomolgus macaques. Squirrel monkeys infected with dengue virus were the reference group. The $P$ value threshold was 0.05.
Squirrel monkeys received similar doses of DENV as cynomolgus macaques from the high exposure group (squirrel monkeys 2.24 [1.98 ; 2.50] $\log_{10}$ PFU, cynomolgus macaques from the high exposure group 1.91 [1.54 ; 2.28], $P$ value 0.15). Squirrel monkeys received lower doses of DENV than ZIKV (ZIKV 3.47 [3.21 ; 3.74], $P$ value = 8.54e-11).

```
 Family: gaussian  ( identity )
Formula:          log_dose ~ group
Data: dose_fixLOD

     AIC      BIC   logLik deviance df.resid
    42.5     49.4    -16.3     32.5       24


Dispersion estimate for gaussian family (sigma^2): 0.18

Conditional model:
                                Estimate Std. Error z value Pr(>|z|)
(Intercept)                       2.2417     0.1341  16.717  < 2e-16 ***
groupCyno.10Mosq.Dengue virus    -0.3340     0.2323  -1.438 0.150466
groupCyno.1Mosq.Dengue virus     -0.9407     0.2509  -3.750 0.000177 ***
groupSquirrel.15Mosq.Zika virus   1.2309     0.1896   6.491 8.54e-11 ***
---
Signif. codes:  0 ‘***’ 0.001 ‘**’ 0.01 ‘*’ 0.05 ‘.’ 0.1 ‘ ’ 1
```

#### S.5 Cytokines - Effect of virus and host species

We assessed, in infected individuals only, the effect of NHP species and virus, for each cytokine separately. To do so, we fitted linear mixed effect models using maximum likelihood, with the $\log_{10}$ cytokine concentration as the response variable, the experiment as the fixed effect: DENV-squirrel was used as the reference, which implied that a significant difference in the DENV-cyno group was a species effect, and a difference in the ZIKV-squirrel group was a virus effect. Day and individual were used as random effects. All concentrations below the limit of detection (LOD) were excluded from analysis initially, for all models. This substantially improved the residuals of

the models, which were checked using the R package DHARMA. To account for multiple testing, we set alpha (the threshold for significance) to 0.001. Results considered significant after this $P$ value correction were compared to corresponding models including measures below LOD (in that case, fixed at LOD). In addition, if an effect of species was significant, the same model was run on control individuals, and we checked through the coefficient estimate and confidence interval whether the species effect in infected individuals was different than in controls. Cytokine concentrations measured before infection and at day 28 post infection were excluded from analyses.
A difference between NHP species during DENV infection was detected in Eotaxin, I-TAC, and RANTES concentrations (Figure S.5). In control individuals, the species difference for these cytokines were of the same magnitude (Figure S.5, Table S.5.4).

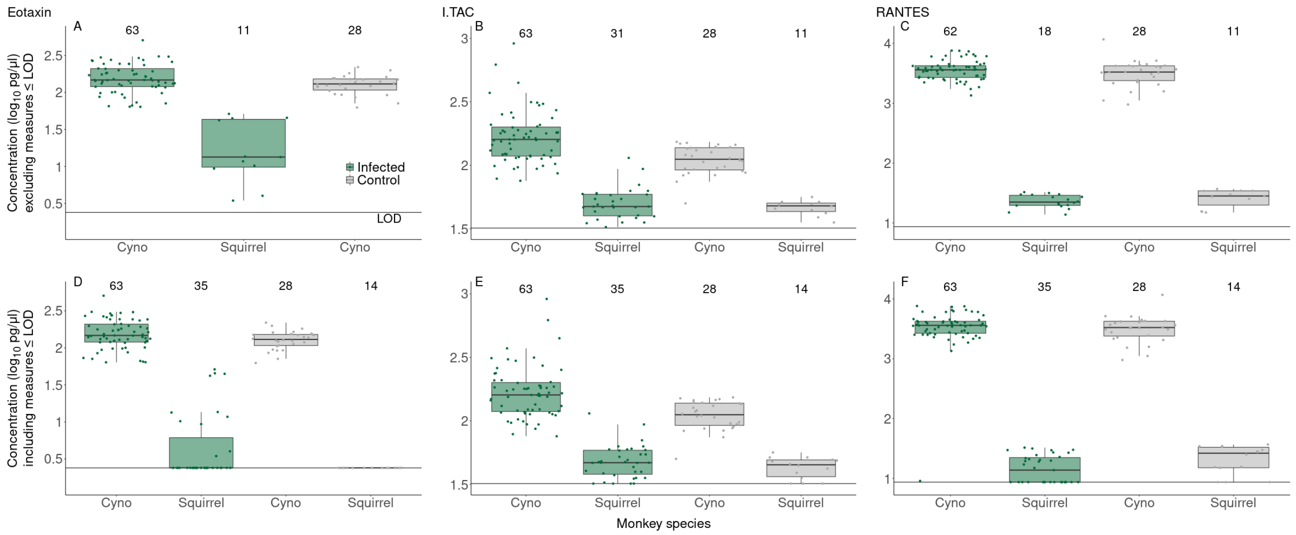


Figure S.5: Effect of monkey species on cytokine concentration, in controls and during DENV infection, for selected cytokines. A-C - Excluding measures below the limit of detection (LOD). D-F - Including measures below LOD (fixed at LOD). Numbers above boxplots indicate the number of datapoints per group. Horizontal black lines show LOD. Scale of y-axis and LOD differ between cytokines, grouped in columns (A,D : Eotaxin, B,E : I-TAC, C,F : RANTES). LOD also slightly differed, for a given cytokine, between monkey species because of the assay. The line shows the minimum LOD for the cytokine.

#### S.6 Individual kinetics of ZIKV replication and transmission

Two monkeys infected with ZIKV, 4683 and 4728, had to be euthanized prior to the end of the experiment following the recommendation of the head veterinarian on staff. On Day 13, NHP 4683 was found during morning rounds hypothermic and hypoglycemic at the bottom of the cage. Shortly after, cluster seizures were observed, and euthanasia was elected. A lesion in the occipital-temporal lobe likely led to the seizures resulting the hypothermia and hypoglycemia. No samples were taken at necropsy to analyze for the presence of ZIKV. Clinical presentations were inconsistent with previous ZIKV reports, but infection causation cannot be ruled out. On Day 15, NHP 4728 had a worsening sore on the back left foot heel that failed to respond for more than 2 weeks to repeated treatments of chlorohexidine and silver sulfadine. These two animals had the lowest weight of all 12 female squirrel monkeys prior to infection; while the mean weight of female squirrel monkeys in this experiment was 676.1g $\pm$ 21g, NHPs 4728 and 4683 weighed 555g and 583g, respectively. These two NHPs also produced the two highest titers of ZIKV of all 10 monkeys infected with the virus (Figure S.6).

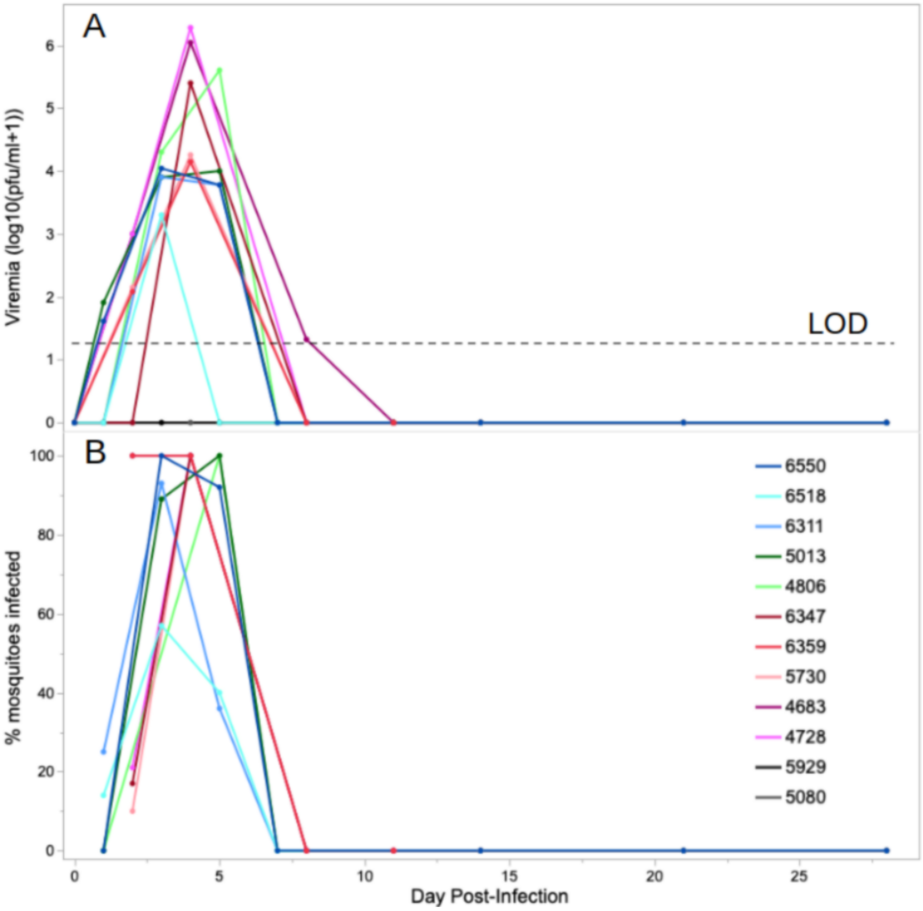


Figure S.6: ZIKV viremia (A) and percentage of mosquitoes infected (B) at designated day post-infection for designated squirrel monkey. Dark lines indicate control animals, blue and purple tone lines indicate cohort 1, and red and orange tone lines indicate cohort 2. Horizontal dashed line shows the limit of detection.

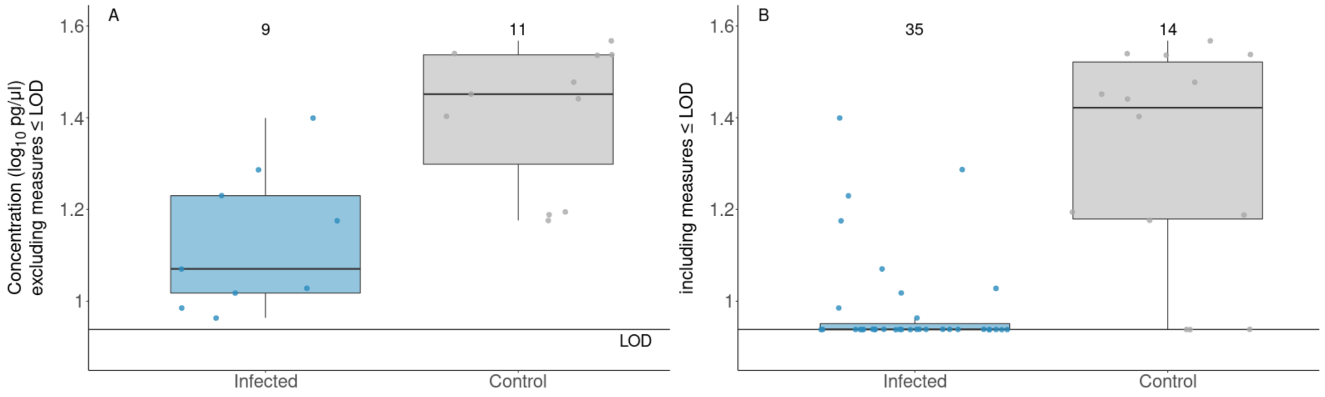


Figure S.7: Effect of Zika virus infection on RANTES concentration. A - Excluding measures below the limit of detection (LOD). B - Including measures below LOD (fixed at LOD). Numbers above boxplots indicate the number of datapoints per group. Horizontal black lines show LOD. See Text S.3.1.

For analyses performed in Sections S.7 and S.8, we corrected some of our initial data to be able to distinguish between likely true absence of viremia and undetectable levels, for the fitting of dose-response relationships. In some cases, non-human primates (NHPs) had no detectable viremia, even after one passage in Vero cells, but did show transmission to mosquitoes (body and/or leg). We assigned a viremia of 10 PFU/ml (half the limit of detection) to these cases. Similarly, if a NHP was not detectably viremic nor transmitted to mosquitoes on a given sampling day, but was viremic and/or transmitted to mosquitoes on the previous and following sampling days, we also assigned a viremia of 10 PFU/ml. Lastly, a cynomolgus macaques which transmitted virus to mosquitoes on day 8 was assigned a viremia of 10 PFU/ml, in the absence of actual viremia measurement. The remaining samples with no detectable viremia even after passage, and no transmission to mosquitoes were considered as true absence of viremia. The resulting data is presented in Figure S.8, and is refered as deduced viremia.

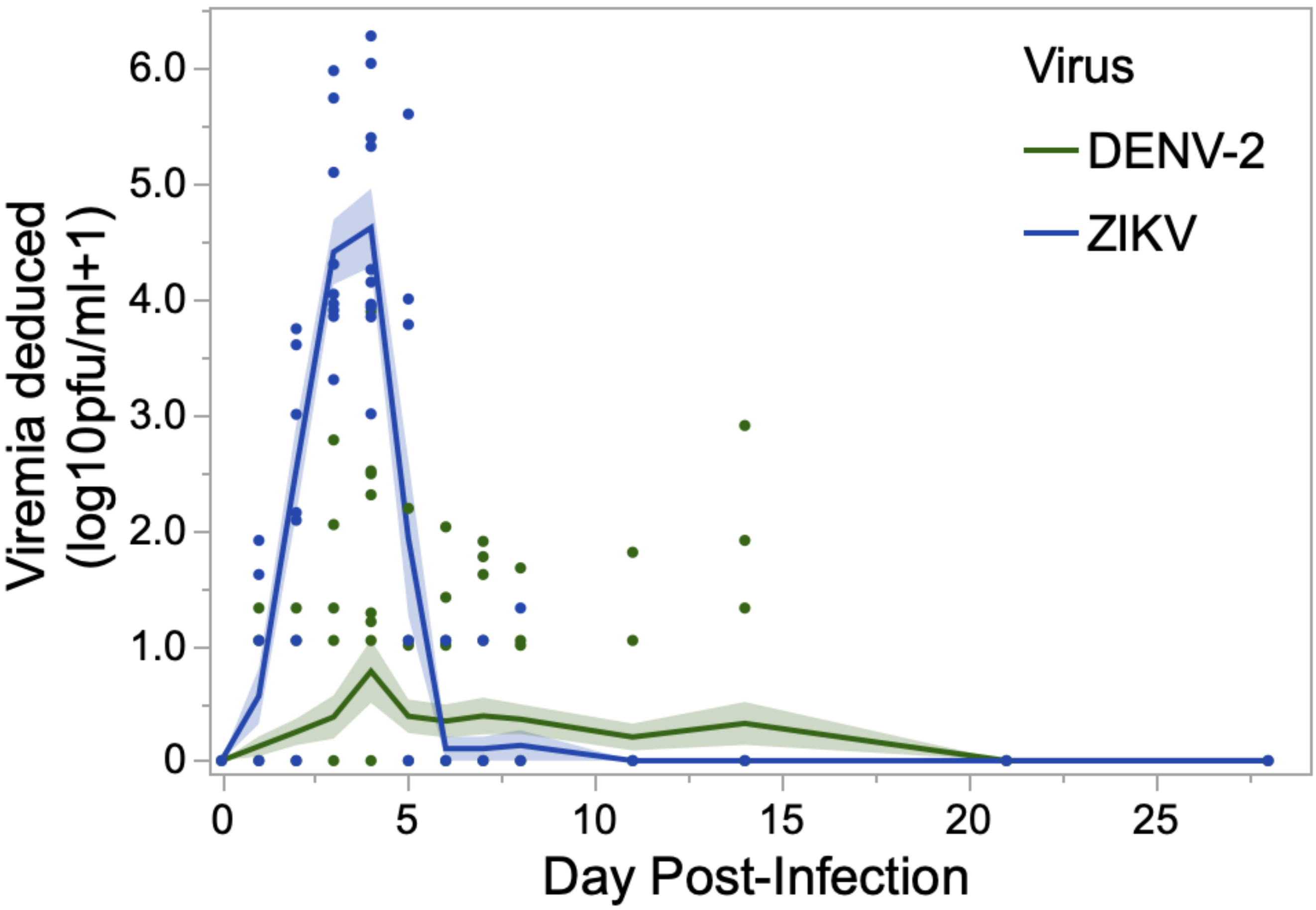


Figure S.8: Deduced viremia from macaques and squirrel monkeys infected with DENV-2 and squirrel monkeys infected with ZIKV at each dpi. Lines show mean and shaded bands show standard errors. See text above for how these viremia values sometimes differ from the initial result of the assays.

#### S.7 Comparison of Dengue and Zika virus transmission from squirrel monkeys to *Aedes albopictus*

To quantify a possible relationship between host infectious titer ($\log_{10}$ PFU/ml) and probability to infect mosquitoes, we first used a flexible fitting approach known as generalized additive model. The probability of mosquito infection was broadly defined, measured by a positive mosquito body or leg. We used a binomial error distribution, and constrained the number of knots (i.e the number of polynomials composing the final curve) to 6. Separate relationships were fitted for dengue and Zika viruses, and transmission from both NHP species was considered for dengue. See Figure 4 in main text.

##### S.7.1 Dengue virus transmission

```
Family: binomial
Link function: logit

Formula:
  cbind(k, N - k) ~ s(log_V, k = 6)

Parametric coefficients:
  Estimate Std. Error z value Pr(>|z|)
(Intercept)   -17.43      37.04  -0.471    0.638

Approximate significance of smooth terms:
  edf Ref.df Chi.sq p-value
s(log_V) 1.841  1.995  4.173   0.118

R-sq.(adj) =  0.409   Deviance explained = 67.4%
UBRE = -0.47875  Scale est. = 1         n = 157
```

##### S.7.2 Zika virus transmission

```
Family: binomial
Link function: logit

Formula:
  cbind(k, N - k) ~ s(log_V, k = 6)

Parametric coefficients:
  Estimate Std. Error z value Pr(>|z|)
(Intercept)  -2.9402     0.8104  -3.628 0.000285 ***
  ---
  Signif. codes:  0 '***' 0.001 '**' 0.01 '*' 0.05 '.' 0.1 ' ' 1

Approximate significance of smooth terms:
  edf Ref.df Chi.sq p-value
s(log_V) 3.232  3.611  68.25  <2e-16 ***
  ---
  Signif. codes:  0 '***' 0.001 '**' 0.01 '*' 0.05 '.' 0.1 ' ' 1

R-sq.(adj) =  0.844   Deviance explained = 85.2%
UBRE = 0.14002  Scale est. = 1         n = 64
```

#### S.8 Comparison of Zika Virus Transmission From Squirrel Monkeys and Dengue Virus Transmission From Humans

We fitted three different functional forms (Eqs. S.1-S.3) to the data, using a maximum likelihood approach. In these equations, $p$ stands for the probability to infect a vector and $V$ is the infectious titer of the host, on a linear scale. Each functional form was fitted with either a binomial likelihood or a beta-binomial likelihood, the latter accounting for overdispersion in the data. We used AICc to select the functional form providing the best fit to data[7]. This fitting procedure was applied twice to obtain separate relationships for mosquitoes' body and leg infection. To not constrain the fitting at the origin, we excluded data considered as true absence of viremia with no transmission to mosquitoes.

$$Logistic : p(V) = \frac{1}{1 + exp(-\beta_1(\log_{10}(V) - \log_{10}(\beta_0)))} \tag{S.1}$$

$$Ferguson : p(V) = 1 - exp(-(\frac{\log_{10}(V)}{\theta_0})^{\theta_1}) \tag{S.2}$$

$$Hill : p(V) = \frac{\log_{10}(V)^{\gamma_1}}{\gamma_0 + \log_{10}(V)^{\gamma_1}} \tag{S.3}$$

Eq. S.1 is the logistic function, with the curve's maximum value fixed at 1 to be on the scale of probabilities. Eq. S.2 [8] was used to study vector competence for dengue when carrying Wolbachia. Eq. S.3, also called the Hill equation, is often used to model biological interactions that demonstrate sigmoidal response, in particular to capture the biomolecular interaction exhibiting cooperativity among two binding molecules[9]. To visualize the uncertainty around fitted curves, we sampled 5000 parameter sets using multivariate normal sampling with the covariance matrix of the fitting procedure.

In the study by Nguyen et al. 2013[10], data came from patients hospitalized at Ho-Chi-Minh city hospital in Vietnam between April and December 2011. DENV-1 cases were associated with the Genotype 1 lineage (different clades), and DENV-2 cases mostly with the Asian 1 lineage (a few Cosmopolitan). Cases with serotypes 3 and 4 were not sequenced. Mosquito infection was measured through the presence of virus in mosquito abdomens. The data we retrieved from the supplementary material of ten Bosch et al. 2018[11] contained 105 data points (28 DENV-1, 13 DENV-2, 16 DENV-3, 48 DENV-4), which is clearly only a subset of the data presented in Figure 2 of [10] (more than 260 data points estimated visually). Note that we did not get an answer from the senior author of [10] when emailed, and that we could not find a valid email address for the first author of the paper.

In the study by Duong et al. 2015[12], viral load and transmission data came from Cambodian participants (Kampong Cham province) infected between June and October of 2012 and 2013. DENV-1 cases were associated with the Genotype 1 lineage, DENV-2 cases with the Asian 1 lineage, and DENV-3 cases with Genotype 1 lineage (we excluded the two DENV-3 datapoints as it was insufficient for fitting). Mosquito infection was measured through the presence of virus in legs and wings. We did not distinguish between classes of disease severity (symptomatic, pre-symptomatic, asymptomatic) in our analyses.

The conversion factors used to transform RNA-emia data into infectious titers were retrieved from Blaney et al. 2005[13], which used strains DENV-1 Nauru/74, DENV-2 Tonga/74, DENV-3 Sleman/78, and DENV-4 Dominica/81. The difference between $\log_{10}$ genome equivalents/ml of serum and $\log_{10}$ PFU/ml of serum were 1.9 for DENV-1, 2.8 for DENV-2, 2.5 for DENV-3, and 1.9 for DENV-4. Because of this conversion, the curve fitted to the DENV-2 dataset from [12] was quite different from the one showed in the initial paper. Indeed, most points were shifted to the left, except for 2 points which corresponded to an absence of detectable viremia with transmission to mosquitoes. Those could not be excluded but were now closer to other points than in the initial RNA-emia scale. This curve was no longer insightful (no sigmoidal shape) and was therefore excluded. All fitting was done on $\log_{10}$(viremia + 1) to accommodate values below 1 PFU/ml.

Tables S.7 and S.8 present the results of model selection and the resulting parameter estimates, for Zika virus only. Figure S.9 presents all dose-response curves relationship fitted along with the data used. Results regarding dose-response relationships using presence of virus in mosquito bodies are presented in the main text.

The best dose-response fit to characterize the relationship between viremia and transmission (measured as infection of the mosquito legs only) of Zika virus by squirrel monkeys was obtained using a logistic equation (Eq. S.1) and a betabinomial likelihood (Figure S.10). This was also the case for DENV-1 data from [12]. For DENV-4 data from [12], the best fit was obtained using the Hill equation Eq. S.3 using a betabinomial likelihood. To compare these curves, we report the estimations of the dose needed to infect the legs of 50% of mosquitoes in Figure S.10.

| Dose-response functional form | Likelihood | Number of parameters | AICc (body) | AICc (leg) |
|---|---|---|---|---|
| Logistic (Eq. S.1) | binomial | 2 | 82.3 | 101.7 |
| | beta-binomial | 3 | **79.6** | **77.5** |
| Ferguson (Eq. S.2) | binomial | 2 | 82.6 | 105.5 |
| | beta-binomial | 3 | 80.5 | 78.9 |
| Hill (Eq. S.3) | binomial | 2 | 82.5 | 113.6 |
| | beta-binomial | 3 | 80.9 | 81.6 |

Table S.7: Model selection for the dose-response relationships predicting the probability of *Aedes albopictus* body or leg infection based on squirrel monkeys' infectious titers. The models with lowest corrected Akaike Information Criterion (AICc, highlighted in bold) are selected.

| Parameter | Vector body infection | | Vector leg infection | |
|---|---|---|---|---|
| | Point-estimate | 95% conf. interval | Point-estimate | 95% conf. interval |
| $\log_{10}(\beta_0)$ | 5.00 | [4.09 ; 5.93] | 2.53 | [1.96 ; 3.13] |
| $\beta_1$ | 0.70 | [0.36 ; 1.04] | 1.12 | [0.59 ; 1.64] |
| overdispersion | 8.58 | [-3.36 ; 20.53] | 2.31 | [-0.02 ; 4.65] |

Table S.8: Parameter estimates for the selected models of Zika virus transmission from squirrel monkeys to *Aedes albopictus*. Model using the logistic function (Eq S.1) fitted with a betabinomial likelihood, accounting for overdispersion in the data. $\log_{10}(\beta_0)$ directly corresponds to the dose ($\log_{10}$) needed to infect the bodies or legs of 50% of mosquitoes.

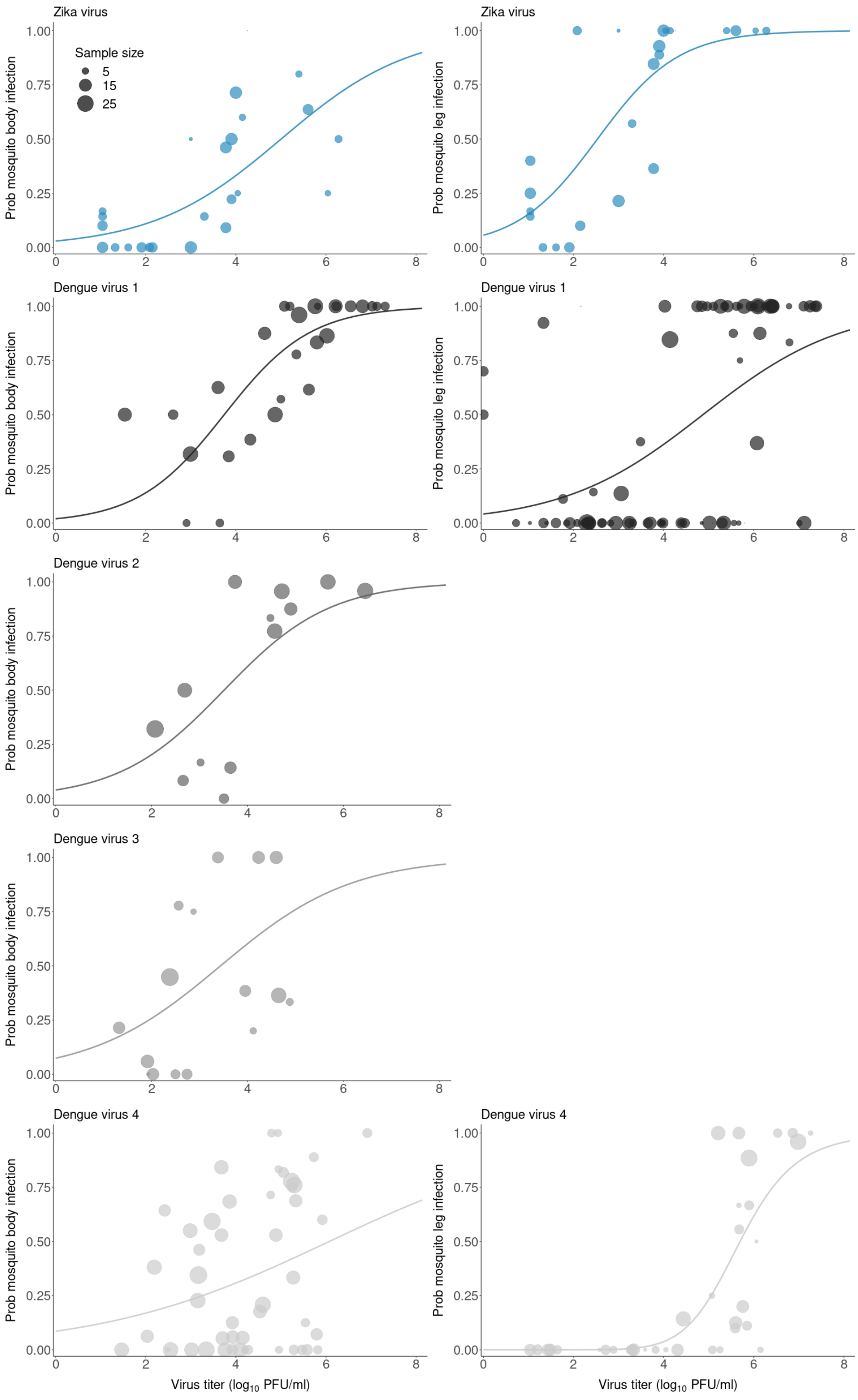




Figure S.9: (Caption next page)

Figure S.9: (Previous page.) Dose-response curves fitted to mosquito body infection data (left column) and mosquito leg infection data (right column). First row is Zika virus transmission data from the present study, next rows are Dengue virus transmission data from the literature (1 row per serotype, data from Nguyen et al. 2013[10] in the left column and from Duong et al. 2015[10] in the right column). Note that as we did not know the value of the limit of detection (LOD) for dengue studies, points with zero viremia and transmission to mosquitoes have been used for fitting, whereas for Zika virus those points were assigned a viremia of 10 PFU (half the LOD of our assays).

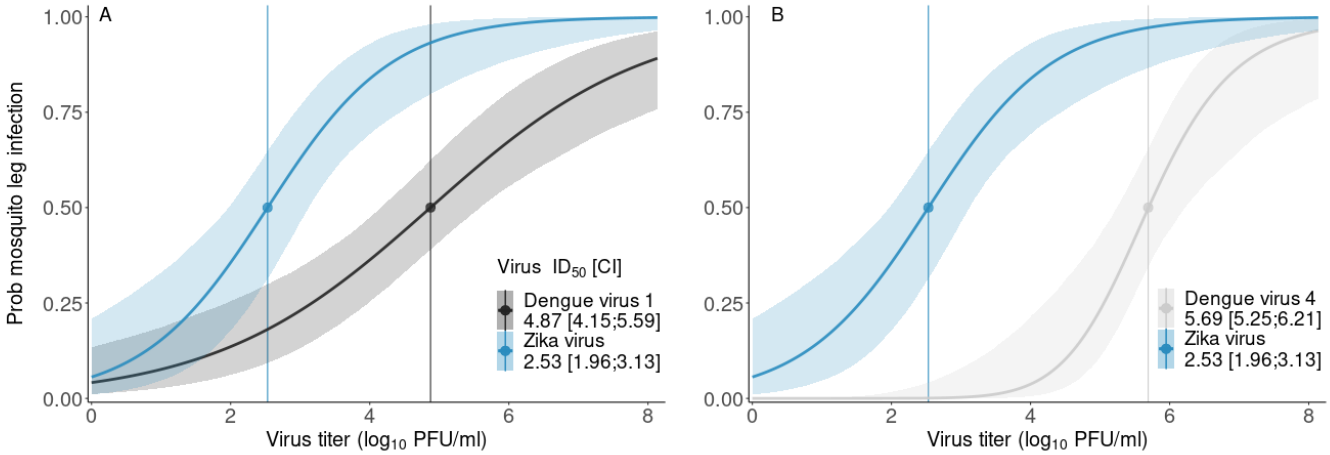


Figure S.10: Relationship between virus titer in serum and transmission of Zika virus to legs of *Ae. albopictus* (blue curves, same curve repeated in each panel) or transmission to legs of *Ae. aegypti* of (A) dengue virus serotype 1, (B) dengue virus serotype 4 (dengue virus data from Duong et al. 2015[10]). Infectious dose 50 ($ID_{50}$), and confidence interval (CI), provided for each designated virus and mosquito species in $\log_{10}$ PFU/ml.

### References

1. Fortman, J., Hewett, T., Bennett, B. Important biologic features. Chapter 1. in *The laboratory nonhuman primate* (eds Fortman, J., Hewett, T., Bennett, B.) pg 17 (Boca Raton, FL: CRC, 2002)

2. Hrapkiewicz K and Medina L. Non-human primates. in *Clinical Laboratory Animal: An Introduction* pg 287 (Blackwell Publishing, Ames, Iowa, USA, 2007)

3. *The Merck Veterinary Manual.* (Whitehouse Station, NJ :Merck & Co., Inc.)

4. Brady, A. G. Research Techniques for the Squirrel Monkey (Saimiri sp.). *ILAR Journal*, **41**, 1 (2000)

5. Marquardt, N. et al. The human NK cell repsonse to Yellow Fever virus 17D is primarily governed by NK cell differentiation independently of NK cell education. *The Journal of Immunology* **195**, 3262–3272 (2015).

6. Björkström, N. K., Strunz, B. & Ljunggren, H.-G. Natural killer cells in antiviral immunity. *Nat Rev Immunol* **22**, 112–123 (2022).

7. Burnham, K. P., Anderson, D. R. *Model Selection and Multimodel Inference : a Practical Information-theoretic Approach.* Second Edition. (Springer Science+Business Media New York, 2002)

8. Ferguson, N. M. et al. Modeling the impact on virus transmission of Wolbachia-mediated blocking of dengue virus infection of Aedes aegypti. *Sci. Transl. Med.* **7**, 279-279ra37 (2015).

9. Somvanshi, P. R., Venkatesh, K.V. Hill Equation. in *Encyclopedia of Systems Biology* (eds Dubitzky, W., Wolkenhauer, O., Cho, K. H., Yokota, H.) 892-895 (Springer New York, 2013)

10. Nguyen, N. M. et al. Host and viral features of human dengue cases shape the population of infected and infectious Aedes aegypti mosquitoes. *Proc. Natl. Acad. Sci. U.S.A.* **110**, 9072–9077 (2013).

11. ten Bosch, Q. A. et al. Contributions from the silent majority dominate dengue virus transmission. *PLOS Pathogens* **14**, e1006965 (2018).

12. Duong, V. et al. Asymptomatic humans transmit dengue virus to mosquitoes. *Proc. Natl. Acad. Sci. U.S.A.* **112**, 14688–14693 (2015).

13. Blaney, J. E., Matro, J. M., Murphy, B. R. & Whitehead, S. S. Recombinant, live-attenuated tetravalent dengue virus vaccine formulations induce a balanced, broad, and protective neutralizing antibody response against each of the four serotypes in rhesus monkeys. *J. Virol.* **79**, 13 (2005).
